# Differentiation marker-negative CD4^+^ T cells persist after yellow fever virus vaccination and contribute to durable memory

**DOI:** 10.1101/2024.03.11.584523

**Authors:** Yi-Gen Pan, Laurent Bartolo, Ruozhang Xu, Bijal Patel, Veronika Zarnitsyna, Laura Su

**Affiliations:** Department of Medicine, Division of Rheumatology, Perelman School of Medicine, Institute for Immunology, University of Pennsylvania, Philadelphia, PA 19104, USA; Corporal Michael J Crescenz VA Medical Center, Philadelphia, PA, 19104, USA; Department of Microbiology and Immunology, Emory University, Atlanta, GA, USA

## Abstract

Factors that contribute to durable immunological memory remain incompletely understood. In our longitudinal analyses of CD4^+^ T cell responses to the yellow fever virus (YFV) vaccine by peptide-MHC tetramers, we unexpectedly found naïve phenotype virus-specific CD4^+^ T cells that persisted months to years after immunization. These Marker negative T cells (T_MN_) lacked CD95, CXCR3, CD11a, and CD49d surface protein expression, distinguishing them from previously discovered stem-cell memory T cells. Functionally, they resembled genuine naïve T cells upon *in vitro* stimulation. Single-cell TCR sequencing detected expanded clonotypes within the T_MN_ subset and identified a shared repertoire with memory and effector T cells. T cells expressing T_MN_-associated TCRs were rare before vaccination, suggesting their expansion following vaccination. Longitudinal tracking of YFV-specific responses over the subsequent years revealed superior stability of the T_MN_ subset and their association with the longevity of the overall population. The identification of these long-lived, antigen-experienced T cells may inform the design of durable T cell-based vaccines and engineered T cell therapies.

## Introduction

Functional immunological memory underlies the protective efficacy of vaccines against subsequent infections (*1, 2*). However, why protection from some vaccines last decades while others wane after a few months remains unknown. A crucial aspect of immune memory involves T cells (*3*). CD8^+^ T cells produce anti-viral cytokines and eliminate infected cells, while CD4^+^ T cells provide key signals for B cell maturation and high-affinity antibody production (*4*). CD4^+^ T cells are also needed to support the expansion and maintenance of functional CD8^+^ T cells and can directly contribute to anti-viral effects (*4–6*). Past studies in mice and humans have identified naïve-like antigen-experienced T cells with superior longevity and plasticity as a source of durable memory (*7–9*). Broadly categorized as stem cell-like memory T cells (T_SCM_), these cells phenotypically resemble naïve T cells by positive CCR7 and CD45RA or negative CD45RO expression, yet they display differentiation markers such as CD95, CXCR3, and CD49d (*9, 10*). In people immunized with the highly efficacious and durable Yellow Fever Virus (YFV) vaccine, class I tetramer analyses identified T_SCM_ as the predominant phenotype of virus-specific CD8^+^ T cells greater than 8 years after vaccination (*10, 11*).

The durability of CD4^+^ T cell memory is less understood. Although capable of differentiating into T_SCM_ cells (*12–15*), CD4^+^ T cells are generally less responsive to homeostatic cytokines IL-7 and IL-15 (*16–18*), which augment T_SCM_ differentiation in cultured CD8^+^ T cells (*19*). Here, we examined virus-specific CD4^+^ T cells after YFV vaccination to delineate key features of durable CD4^+^ T cell responses. YFV-specific CD4^+^ T cells were identified and tracked longitudinally by direct *ex vivo* class II peptide-MHC (pMHC) tetramers staining. YFV-specific CD4^+^ T cells existed in various memory states, including a T_SCM_ subset. Unexpectedly, some tetramer-labeled T cells remained negative for all measured differentiation markers several months after vaccination.

Focused analyses of these marker-negative T cells (T_MN_) showed that they responded to cognate peptide stimulation *in vitro* and had likely undergone proliferation *in vivo* by clonal expansion. Further, their T cell receptor (TCR) sequences overlapped with that of classically defined memory T cells of the same specificity, implicating a shared clonal origin. T_MN_ cells functionally resembled genuine naïve T cells, exhibited superior stability over other memory subsets, and were associated with the long-term persistence of the overall population years after vaccination. Our findings expand the current definition of antigen-experienced T cells to include those that retain an undifferentiated phenotype. These cells opened new avenues for understanding the durability of immune responses and developing strategies to enhance long-lasting immunologic memory.

## Results

### Detection of naïve-like CD4^+^ T cells after YFV vaccination

We had previously performed a longitudinal study of YFV-specific CD4^+^ T cells to evaluate the impact of pre-existing repertoire on T cell responses to primary immunization with the YFV vaccine (*20*). Starting with this dataset, we examined the features of memory T cells that developed at least 7 months after vaccination. This showed that approximately half of the YFV-specific memory pool consisted of central memory T cells (T_CM_), with about 21% of tetramer^+^ cells retaining a naïve-like CD45RO^-^CCR7^+^ phenotype (Fig. 1A-B). Proportionally, the abundance of CD45RO^-^CCR7^+^ subset was highest before vaccination, decreased initially post-vaccination, then reaccumulated several months later (Fig. 1C). By frequency, CD45RO^-^CCR7^+^ tetramer^+^ cells increased steadily after vaccination (Fig. 1D). The frequency of CD45RO^-^CCR7^+^ T cell subset did not differ by donor age but was instead associated with the robustness of the response (Fig. S1A-C). CD45RO^-^CCR7^+^ YFV-specific T cells were more abundant in populations that reached a higher frequency and positively correlated with the fold-change between the peak and the pre-vaccine baseline (Fig S1B-C). At a later memory time point, the frequencies of CD45RO^-^CCR7^+^ YFV-specific T cells were higher within larger populations that were recruited into the memory pool (Fig. 1E-F). These data suggest that CD45RO^-^CCR7^+^CD4^+^ T cells is a feature of an effective T cell response.

**Figure 1:**
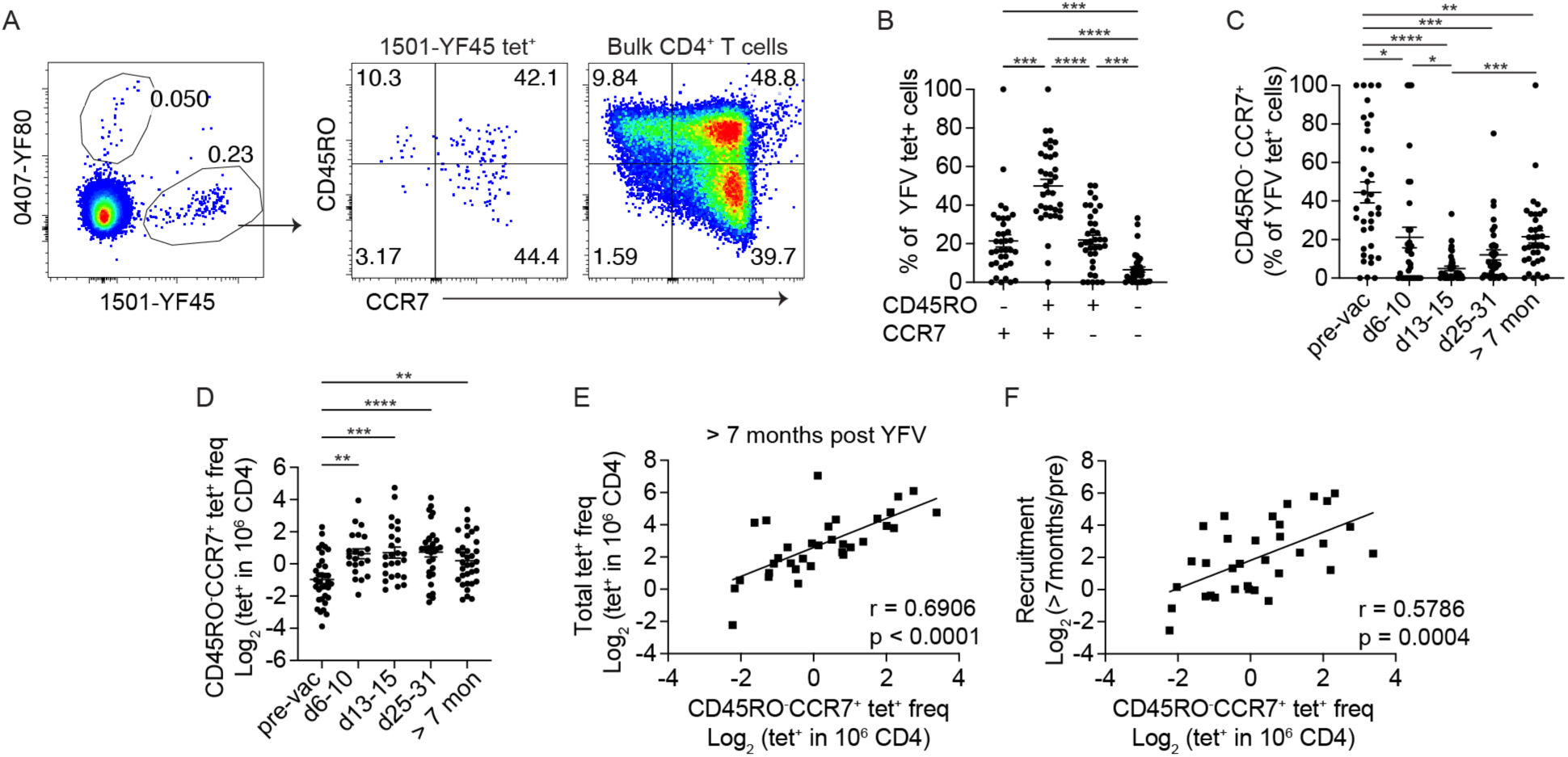
Identification of a CD45RO^-^CCR7^+^ subset of virus-specific CD4^+^ T cells after YFV vaccination. (A) Direct *ex vivo* tetramer and antibody staining of a representative YFV tetramer^+^ population using blood collected about 7 months after YFV vaccination. (B) The percentage of YFV tetramer^+^ T cells with the indicated combination of CD45RO and CCR7 expression. Plot summarizes data from 36 specificities 7 to 34 months after YFV vaccination from 7 donors. (C-D) The abundance of CD45RO^-^CCR7^+^ YFV tetramer^+^ CD4^+^ T cells in 7 healthy subjects was quantified as a percentage of tetramer^+^ cells (C) or by frequency (D). Each symbol represents data from a distinct YFV-specific population. Experiments were repeated an average of 3.3 times. (E-F) Correlation between the frequency of CD45RO^-^CCR7^+^ subset with the corresponding overall frequency (E) and the fold change between memory frequency and the pre-vaccine baseline (F) (n = 36). RM one-way ANOVA (B) or Mixed-effect analysis (C and D) was performed and corrected with Tukey’s multiple comparisons test. For E and F, Pearson correlation was computed.

### Post-immune T cells are heterogeneous and include a differentiation marker negative subset

We hypothesized that the post-vaccine CD45RO^-^CCR7^+^ subset largely consisted of T_SCM_ cells as in CD8^+^ T cells (*10, 11*). To test this, we performed tetramer staining on 28 YFV-specific CD4^+^ populations from 7 individuals, recognizing 16 unique epitopes with antibodies against T_SCM_-associated markers, CXCR3, CD95, CD11a, and CD49d (Tables S1 and S2). Staining with this broader antibody panel on blood collected 7 to 48 months after vaccination indeed identified CD45RO^-^CCR7^+^ tetramer^+^ T cells that expressed one or more T_SCM_ markers. However, we noted that a portion of CD45RO^-^CCR7^+^ CD4^+^ T cells remained negative for CXCR3, CD95, CD11a, and CD49d expression (Fig. 2A, S2A-B). To gain further insights into the heterogeneity within the CD45RO-CCR7^+^ subset, we combined 1465 YFV-specific CD4^+^ T cells from one donor and visualized combinatorial antibody staining on UMAP using the Spectre pipeline (*21*). This identified regions with low CD45RO and high CCR7 signals, which encompassed a CXCR3^+^ (cluster 0) and a T_SCM_ marker negative population (cluster 4) (Fig. 2B-D). We defined CD45RO^-^CCR7^+^ cells lacking any measured differentiation markers as marker-negative T cells (T_MN_) and classified those expressing at least one of CXCR3, CD95, CD11a, or CD49d as T_SCM_ cells (Table S3). On average, a quarter of the CD45RO^-^CCR7^+^ subset consisted of T_MN_ cells (Fig. 2E). Among T_SCM_ cells, the majority expressed CXCR3 alone or in combination with other differentiation markers (Fig. 2E-F, S2C). Finding antigen-specific T cells that do not express known memory or T_SCM_ markers after a clear prior exposure was unexpected. To test if T_MN_ cells functionally behave like antigen-experienced T cells despite lacking surface markers of differentiation, we treated post-vaccine PBMCs with PMA and ionomycin for 4 to 5 hours. Antigen-specific T cells were captured by tetramers, divided into distinct phenotypic subsets, and analyzed for TNF-α and IFN-γ production. This showed that post-immune T_MN_ subset produced significantly less cytokines compared to memory T cells within the same tetramer^+^ population (Fig. 2G-H). Thus, YFV vaccination induced a diverse post-immune repertoire that included CD4^+^ T_SCM_ cells and a naïve-like T_MN_ population that lacked phenotypic and functional features of antigen experience.

**Figure 2:**
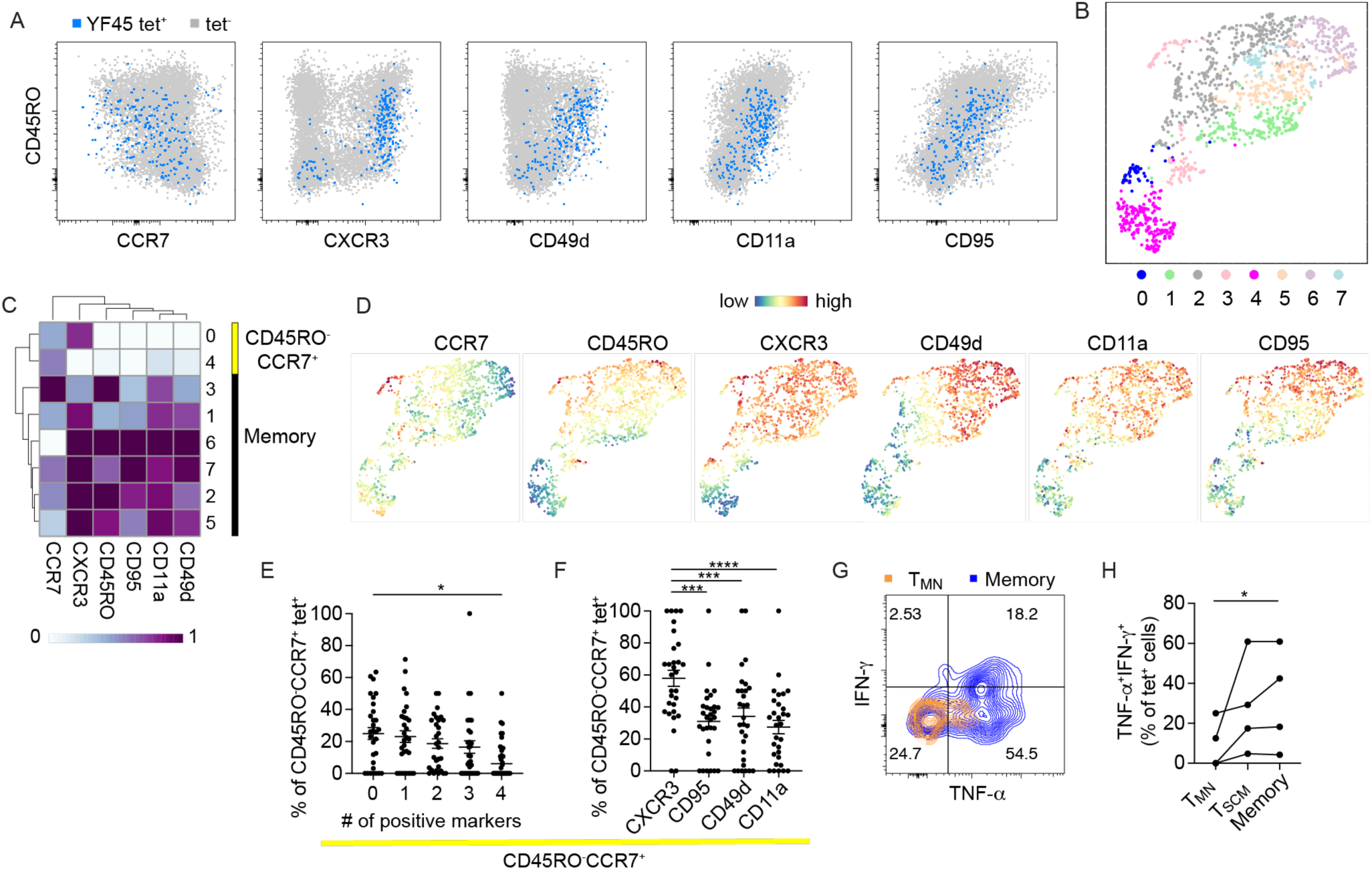
Post-vaccine CD4^+^ T cells are heterogeneous and include naïve-like subsets. (A) FACS plots show the expression of the indicated marker on a representative YFV-specific population. The tetramer^+^ population is overlaid onto tetramer^-^ bulk CD4^+^ T cells. (B) UMAP displays Phenograph-defined clusters. Data combine 1465 CD4^+^ cells labeled by 7 YFV tetramers from HD3. (C-D) The staining intensity of individual markers is shown on a heatmap for each cluster (C) or displayed on the UMAP. (E-F) The relative abundance of CD45RO^-^CCR7^+^ YFV-specific T cells by the indicated numbers of markers (E) or the type of markers (F). Frequency in F combines all cells positive for a particular marker within the CD45RO^-^CCR7^+^ subset. Each symbol represents a tetramer^+^ population (n = 28). Experiments were repeated an average of 2.5 times. (G) PBMCs were stimulated for 4 - 5 hours by PMA and ionomycin and assayed for cytokine production by intracellular cytokine staining. The plot shows representative TNF-a and IFN-g expression by T_MN_ cells and non-CD45RO^-^CD28^+^ (memory) T cells from the same tetramer-labeled population. (H) T cell responses by TNF-a and IFN-g production for the indicated phenotypic subset. Each population was identified with a pool of 5-7 tetramers of the same DR allele, using cells from 3 donors. For E and F, RM one-way ANOVA was performed and corrected with Tukey’s multiple comparison test. For H, the Friedman test was performed and corrected using Dunn’s multiple comparison test.

### Virus-specific T_MN_ cells respond to antigens

We were intrigued by the existence of virus-specific T cells that retained a naïve functional phenotype after vaccination. Past studies have identified non-stimulatory TCR interactions that decoupled T cell activation from ligand binding (*22*). The impaired ability to respond productively to antigens may be one reason why some tetramer-labeled T cells retained a naïve phenotype. To investigate this possibility, we quantified T_MN_, T_SCM_, and T_CM_ cells for differences in their functional avidity by peptide stimulation. YFV-specific T cell clones were generated using samples from two donors obtained 7 to 8 months after YFV vaccination. Among the 48 clones that grew, 40 clones (90%) had the correct specificity by tetramer re-staining and/or response to peptides (Fig. 3A-B, S3A). We did not identify peptide-nonresponsive T cells as all clones that were stained with tetramers responded to peptide stimulation. To determine if T_MN_ cells might be harder to activate due to a lower functional avidity, we divided the clones according to their direct *ex vivo* phenotype and selected 5 clones each from T_CM_, T_SCM_, T_MN_ groups for further analyses. YFV-specific clones were stimulated with decreasing concentrations of the cognate peptide and analyzed for response by cytokine production (Fig. S3B). T_MN_-derived clones responded similarly to peptides by TNF-α production, with no significant differences in maximal effective peptide concentration (EC50) values between groups (Fig. 3C-D). T cell clones, regardless of their *ex vivo* phenotypes, also produced similar levels of IFN-γ, IL-2, and had comparable TNF-α^+^IFN-γ^+^IL-2^+^ co-expression (Fig. 3E). In addition, we evaluated the proliferative capacity of T_MN_, T_SCM_, and T_CM_-derived clones by CellTrace Violet (CTV) dilution and observed no significant differences in the proliferative response to peptide stimulation (Fig. 3F-G, Fig. S3C). Thus, TCR-ligand engagement is likely intact for vaccine-specific T cells that retained a naïve phenotype after vaccination.

**Figure 3:**
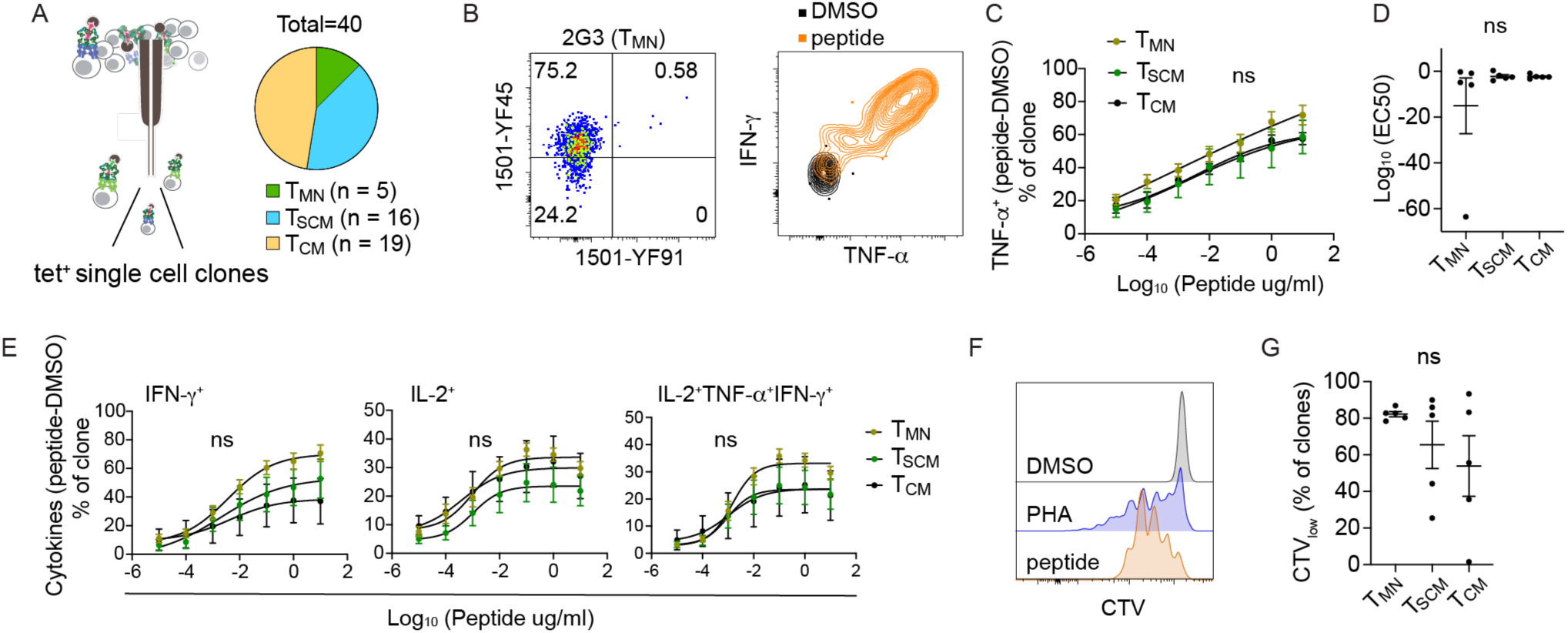
T_M_N-derived T cell clones respond to antigen stimulation. (A) Schematics of single-cell T cell cloning. Post-vaccine T cells from HD2 and HD3 were stained with 1501-YF45 tetramers, sorted based on T_MN_, T_SCM_, or T_CM_ phenotypes, and expanded for 2 to 3 weeks in culture. (B) *In vitro* expanded T cell clones were restained with tetramers and cultured with vehicle or peptide-treated monocyte-derived dendritic cells. Representative plots show tetramer staining and cytokine production by intracellular cytokine staining. (C-D) T cell clones were stimulated with decreasing concentrations of YFV peptide. The response was measured by TNF-a production (C) and quantified by EC50 values after subtracting the background signal from vehicle-treated control (D). (E) Peptide dose response of T cell clones by IFN-g, IL-2, and IL2^+^TNF-a^+^IFN-g ^+^ production. (F) Representative histograms show CTV dilution in response to 10ug/ml of peptide stimulation. (G) Plot summarizes the frequency of CTV_low_ population after a 5-day culture for clones in each phenotypic group. All experiments were repeated at least twice with n= 5 in each group. For (C) and (E), RM two-way ANOVA was performed and corrected with Tukey’s multiple comparison test. For (D) and (G), Kruskal-Wallis test and Dunn’s multiple comparison test were used.

### T_MN_ cells are clonally related to memory and effector T cells

While T_MN_ cells respond well to antigens *in vitro*, it remains possible for them to be less competitive in resource-limiting environments. To investigate this, we reason that we can use TCR sequences to infer stimulation and proliferative response *in vivo*. Because T cell progenies originating from a T cell express identical TCR sequences, we can further leverage these sequences as molecular barcodes to investigate the clonal relationship between distinct phenotypic subsets. However, capturing sufficient numbers of T_MN_ cells was challenging due to their limited number within the available blood samples. To overcome this problem, we generated new tetramers using affinity-matured DR monomers containing mutations that enhanced CD4 binding to improve the overall capture efficiency (*23*). When compared to the wild-type (wt) DR, these tetramers stained a larger population of T cells without significantly skewing the phenotypic proportions (Fig. S4A-C). In total, we sorted single cells from 5 tetramer-labeled populations and obtained TCR sequences from 607 YFV-specific CD4^+^ T cells after amplification and sequencing (Fig. 4A, Table S4). Consistent with clonal expansion after vaccination, over 70% of the sequences were identified in more than one tetramer-labeled T cell. Among expanded sequences, 25 to 52% were abundant and found in at least 10 individual T cells (Fig. 4B). Most T cells displayed a T_CM_ or T_EM_ phenotype based on antibody staining at the time of sorting. T_MN_ phenotype was infrequent, expressed by 3 to 4% of sequenced T cells and confined to the two most extensively sequenced populations recognizing YF45. Consistent with *in vivo* expansion, T_MN_ cells did not preferentially express unique TCRs, but rather, they were distributed across various clone sizes (Fig. 4C). We focused the subsequent analyses on YF45-specific T cells that included the T_MN_ subset. Early post-vaccine measurements of YF45-specific T cells from HD2 and HD3 showed that both populations had generated robust responses to the YFV vaccine (Fig. 4D) (*20*). In agreement with an antigen-driven response, T_MN_ cells contained expanded clonotypes and shared overlapping sequences with various memory subsets (Fig. 4E-F, S4E). In separately generated T cell clones from the same individuals, T_MN_-derived clones expressed TCRs that matched the sequences from *ex vivo* sorted T cells of diverse clone sizes and phenotypes (Fig. S4F).

**Figure 4:**
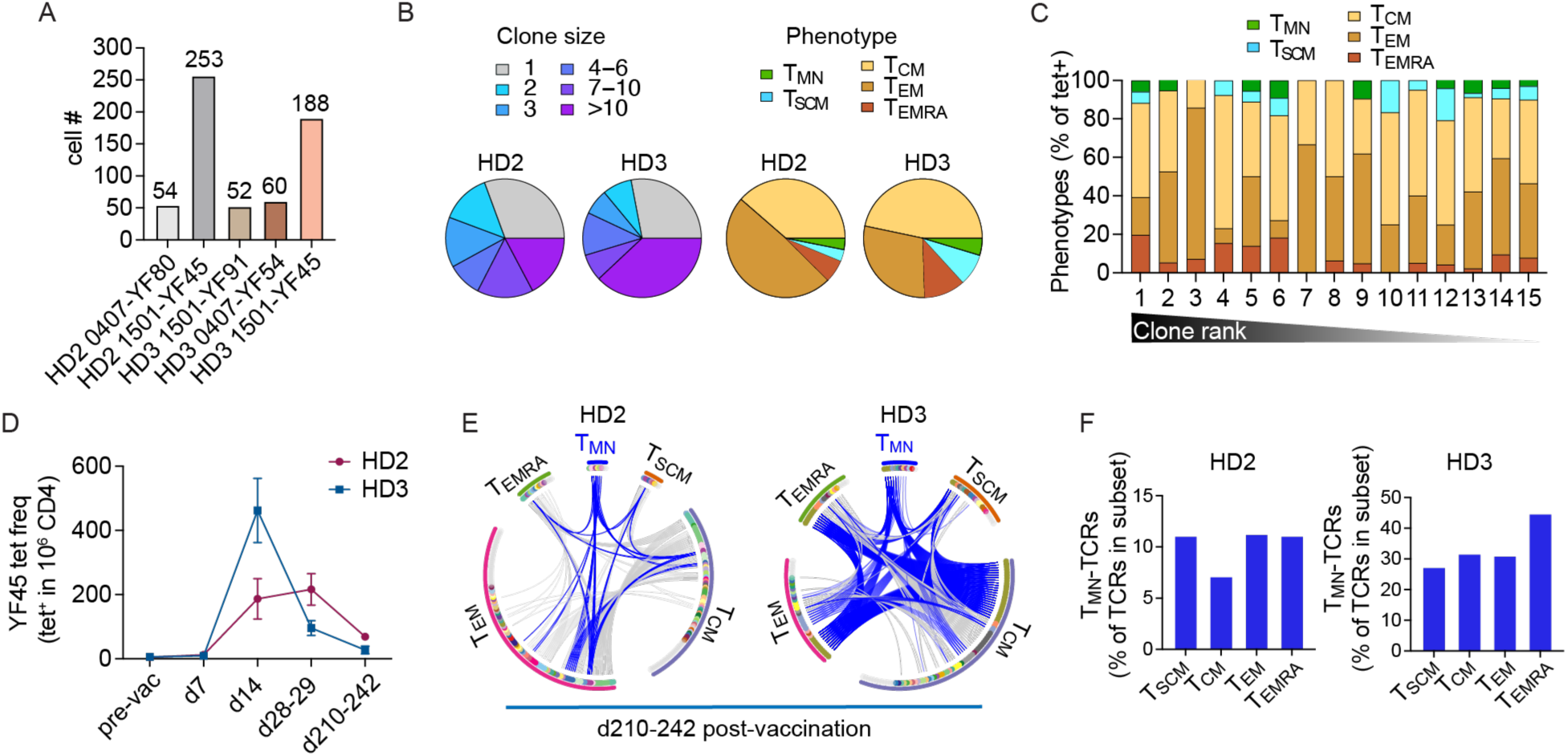
T_M_N cells are clonally related to memory T cells. (A) The plot summarizes the number of cells sequenced from indicated specificity and donor collected 242 (HD2) and 210 (HD3) days after primary YFV vaccination. (B) Clone size and phenotypic distribution of YFV tetramer^+^ populations in A. Phenotypic information were obtained by index sorting. T_MN_ (CD45RO^-^CCR7^+^CXCR3^-^CD95^-^CD11a^-^CD49d^-^), T_SCM_ (CD45RO^-^CCR7^+^ and positive for at least one of CXCR3, CD95, CD11a, or CD49d), T_CM_ (CD45RO^+^CCR7^+^), T_EM_ (CD45RO^+^CCR7^-^), T_EMRA_ (CD45RO^-^CCR7^-^). Cells with ambiguous phenotypes were excluded. (C) Distribution of phenotypes in B by clonotype frequency, ranked from largest to unique clonotypes. (D) Pre-vaccine frequency and early post-vaccine dynamics of YF45-specific T cells preceding the memory time point. (E) Each circus plot represents TCRs from YF45 tetramer^+^ cells obtained 242 (HD2) or 210 (HD3) days after vaccination, separated by the associated indexed phenotypes. Cells are ordered by frequency within each arc. Gray marks cells expressing unique TCRs, other colors represent expanded or shared sequences. Shared TCRb or TCRa/b, when a TCRa is available, is connected by a line across distinct phenotypic subsets. Blue lines highlight TCRs from T_MN_ cells that are shared with cells expressing other phenotypes. (F) The percentage of TCRs in each memory subset that matched T_MN_-derived sequences.

The presence of shared TCR sequences with memory T cells, together with clonal expansion, suggest that T_MN_ cells had encountered and responded to antigens. Alternatively, there could be numerous naïve T cells in the precursor repertoire, some of which could retain a naïve phenotype if only a subset was recruited into the vaccine response. To investigate this possibility, we examined the pre-vaccination repertoire of YF45-specific T cells in these individuals to determine if T_MN_-associated TCRs were abundant before vaccination (*20*). The changes in clonal dynamics were assessed by tetramer staining, sorting, and sequencing the TCRs of YF45-specific T cells from blood collected 14 days after vaccination. In total, we examined TCR sequences from 129 precursor T cells and 238 effector T cells (Fig. 5A, Table S5). Before vaccination, no pre-vaccine TCRs matched T_MN_-derived TCRs from HD2 and only one sequence was identified in HD3. This shared TCR mapped to a unique sequence and not to the expanded pre-existing clonotypes in this individual. By contrast, 7% (HD2, 5 cells) and 23% (HD3, 38 cells) of TCRs in day 14 blood samples expressed a T_MN_-associated TCR (Fig. 5B). Matched T cells in the day 14 sample expressed a variety of differentiation phenotypes and included expanded clonotypes (Fig. 5C). Together, these data indicate that T_MN_ precursors were rare in the pre-immune repertoire and underwent expansion in response to antigen stimulation after vaccination.

**Figure 5:**
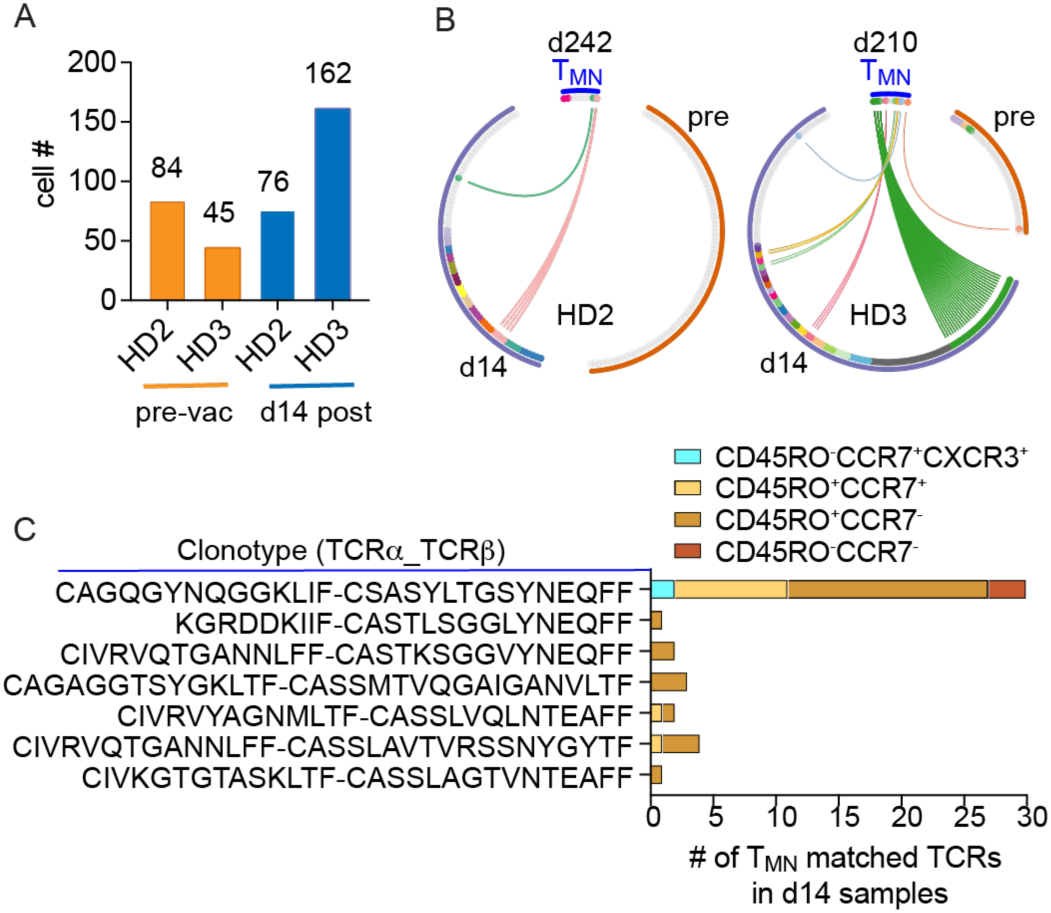
T_M_N cells are clonally related to effector T cells. (A) The number of TCRs from YF45 tetramer^+^ T cells from the indicated donors, before vaccination and 14 days after YFV vaccination. (B) Lines link T_MN_-derived TCRs in the day 210-242 post-vaccine samples with matched TCRs expressed by T cells in a previous time point from each donor. Shared TCRs are matched by TCRb or TCRa/b when a TCRa is available. (C) The CDR3 sequences and the phenotypes of T cells in the d14 sample that matched a T_MN_-derived clonotype in the memory time point.

### T_MN_ cells contribute to durable memory

While memory T cells are essential for generating rapid recall responses, naïve T cells are known for their persistence (*24, 25*). We hypothesize that this unique naïve-appearing antigen-experienced subset would retain this key property and contribute to durable immune responses. To test this idea, we analyzed additional time points from five donors who had longitudinal PBMCs collections up to 6.7 years after YFV vaccination (Fig. 6A, Fig. S5A). Past modeling of cellular turnover suggests that different phenotypic subpopulations undergo separate and distinct *in vivo* dynamics (*26, 27*). To evaluate the stability of individual phenotypic subsets, we subdivided 19 YFV-specific populations according to T_MN_, T_SCM_, T_CM_, T_EM_, and T_EMRA_ phenotypes based on CD45RO, CCR7, CD95, CXCR3, CD11a, and CD49d expression. Their time-dependent change was quantified as a fitted slope using a mixed-effects exponential decay model. This revealed different rates of decay between cells in distinct differentiation states. CD4^+^ T_EM_ cells had the largest negative slope, indicating the greatest decrease over time. In contrast, T_MN_ cells exhibited remarkable stability, with no discernible decline observed during the follow-up period. The stability of the T_MN_ subset significantly surpassed that of other phenotypic subsets, including T_SCM_ and T_CM_ cells, which are typically considered to be long-lived (Fig.6B).

**Figure 6:**
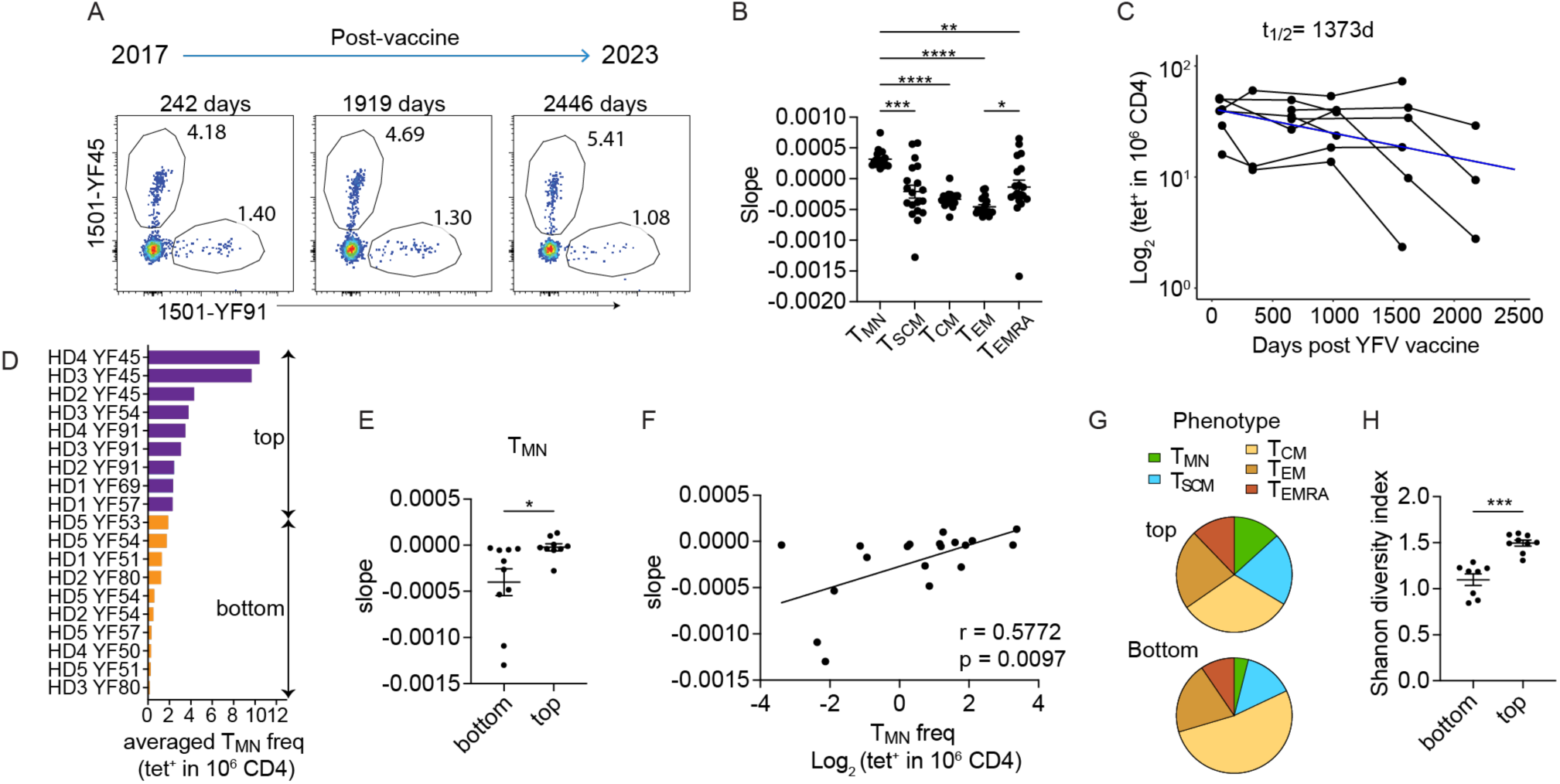
T_M_N cells are stable and associated with durable T cell memory. (A) Representative plots show YFV-specific CD4^+^ T cells over the indicated time points from HD2. (B) Each tetramer^+^ population of a given specificity was subdivided according to phenotypes. The change over time for each phenotypic subset was quantified by the estimated slope using a mixed effects exponential decay model (n = 19 tetramer^+^ populations from 5 donors). (C) A mixed effects exponential decay model fitted to the dynamics of YFV-specific CD4^+^ T cells after a single YFV vaccination (n = 8 populations, combined from donors 4 and 5). The estimated decay (blue line) was used for calculating the half-life (t_1/2_). (D) Ranking of tetramer^+^ populations by the averaged frequency of T_MN_ T cells within each population across all time points. (E) Plot summarizes the estimated slopes of individual tetramer^+^ populations, divided into top and bottom halves by T_MN_ frequency in D. (F) The correlation between slopes characterizing the change over time for the overall tetramer^+^ populations and their corresponding averaged T_MN_ frequencies. (G) Pie-charts show the distribution of memory subsets. Populations were divided into top and bottom groups by the first measured T_MN_ frequency obtained within 1-2 years after YFV vaccination. (H) Phenotypic diversity of each tetramer^+^ population was quantified using Shannon Diversity Index, categorized into top or bottom groups based on T_MN_ frequency as in G. Each symbol represents one tetramer^+^ population. Experiments were repeated an average of 2.3 times. Data are represented as mean ± SEM. For (B), RM one-way ANOVA was performed and corrected with Tukey’s multiple comparisons test. Welch’s t-test was performed for (E) and (H). For (F), Spearman correlation was performed.

Next, we examined the decay kinetics of the overall YFV-specific CD4^+^ T cell responses. Because some data were generated before switching to modified DR, paired analyses by wildtype and modified tetramers on the same blood sample were used to generate an equation for normalizing the frequencies across experiments (Fig. S4D). Among the five donors followed longitudinally, two received one YFV dose as typical for the YFV vaccine, while three had been revaccinated 7 months to a year after the initial dose (Table S6). We grouped the donors based on vaccine dosing to model the frequency of each tetramer^+^ population over time. This revealed a highly durable CD4^+^ T cell memory response after a single dose that becomes further stabilized after re-vaccination (Fig. S5B). Consistent with the longevity of YFV vaccine-mediated protection, YFV-specific CD4^+^ T cells displayed an average half-life (t_1/2_) of close to 4 years after one YFV immunization (Fig. 6C). We observed more T_MN_ cells in the two-dose group, although the difference was not statistically significant (Fig. S5C). Considering the heterogeneity at the population level, we analyzed the tetramer^+^ populations by T_MN_ frequency and divided them into top and bottom halves (Fig. 6D). This showed that populations with more T_MN_ cells were more stable compared to populations in the bottom T_MN_ group (Fig. 6E). The T_MN_ frequency within a given virus-specific population also demonstrated a positive correlation with the stability of the overall population (Fig. 6F). By contrast, we did not find significant differences between high and low groups based on T_SCM_, T_CM_, T_EM_, and T_EMRA_ frequencies (Fig. S5D). On the phenotypic level, all tetramer^+^ populations contained various memory subsets, but the top T_MN_ group was more phenotypically diverse. We divided populations based on the first T_MN_ frequency obtained within the 1-2 years after YFV vaccination and showed that those having more T_MN_ cells exhibited a higher diversity of differentiation states over time as measured by the Shannon diversity index (Fig. 6G-H). Collectively, these data highlight the stability of the T_MN_ subset and uncover their association with durable and diverse T cell memory after YFV vaccination.

## Discussion

We examined CD4^+^ T cell memory to YFV vaccination to define key features of durable memory by direct *ex vivo* class II tetramer staining and enrichment. This showed a diverse memory pool comprised of various differentiation states after YFV vaccination. A portion of YFV-specific CD4^+^ T cells acquired T_SCM_ phenotype after vaccination as in CD8^+^ T cells. Unexpectedly, we also uncovered antigen-experienced T cells that lacked typical markers of T_SCM_ and other memory cells, including CD95, CD11a, CD49d, and CXCR3. Similar to genuine naïve T cells, T_MN_ cells expressed the lymphoid homing chemokine receptor, CCR7 (*28*), and functionally resembled naïve T cells after polyclonal stimulation by PMN and ionomycin. Despite their naïve appearance, T_MN_ cells are antigen-experienced. T_MN_ cells engaged with and responded to antigens *in vitro*. T_MN_ responses to the YFV vaccine *in vivo* were supported by single-cell TCR sequencing, which revealed clonal expansion and a shared repertoire with memory T cells. Longitudinal clonal tracking further identified an expansion of T_MN_-associated TCRs during effector response. Thus, our analyses of CD4^+^ T cell responses to the highly durable and efficacious YFV vaccine revealed that some human CD4^+^ T cells can appear indistinguishable from genuine naïve T cells despite prior antigen experience and expansion.

A T cell is typically referred to as naïve if it has not yet encountered its specific cognate antigen(s). Because a clear antigenic history is often not available, especially in human studies, specific surface markers are commonly used to infer antigen experience (*29, 30*). However, with the advances in single-cell technologies, there is increasing appreciation for the complexity within the naïve compartment. A recent multi-omic analysis discovered age-related epigenetic and transcriptional changes in naïve CD4^+^ and CD8^+^ T cells (*31*). Naïve T cells were defined by CD45RA, CCR7, and CD27 expression using multiple sequencing modalities. Even when naive T cells were further characterized by the lack of CD49d, CD95, and IFN-γ expression, these cells showed distinct chromatin accessibility and transcription factor expression between children and older adults (*31*). The TCR repertoire also undergoes age-related changes and displays a decrease in repertoire diversity in the elderly (*32*). Generally, these changes are thought to have occurred by homeostatic mechanisms that maintain the peripheral naïve repertoire (*33, 34*). While we did not investigate how cytokines impact T_MN_ differentiation, cells uniquely driven by a cytokine-mediated bystander response would not be expected to have a similar TCR repertoire as memory and effector T cells. Partial recruitment of precursor T cells remains formally possible, although T_MN_ precursors were not numerous and past studies in mice suggest a highly efficient recruitment process (*35, 36*). For CD8^+^ T cells expressing OT-1 transgenic TCR, even the weakest altered peptide ligands, about 700-fold less potent than the wt sequence, induced effector response and generated memory T cells (*36*). Regardless of the mechanism of T_MN_ differentiation, our data indicate that some cells considered naïve by phenotypic criteria have actually encountered and responded to foreign antigens. Over a lifetime, these cells may accumulate and add to non-antigen experienced T cells as a part of the naïve repertoire, thereby changing naïve T cell composition over time.

Only a few select vaccines are capable of mediating life-long protection. How durable immunological memory is maintained remains a key unresolved question. While Memory T cells are the cornerstone of protective immunity by virtue of their ability to rapidly initiate a functional response to pathogen rechallenge, naïve T cells possess superior self-renewal capacity and differentiation plasticity (*3, 24, 37*). Considering the phenotypic and functional similarities between T_MN_ and naïve T cells, we asked if T_MN_ cells contribute to the longevity of T cell response after YFV vaccination. Our findings revealed remarkable stability of T_MN_ cells, exhibiting minimal decay for nearly 7 years. Their ability to persist suggests that T_MN_ cells could potentially support the longevity of the overall immune response, extending it beyond the lifespan of individual memory T cells. Consistent with this model, T_MN_ cells are more abundant in durable CD4^+^ populations that are stable over time. Based on the diverse memory phenotypes in T_MN_-enriched populations, we further speculate that T_MN_ cells have the potential to differentiate into multiple states, thereby contributing to the phenotypic diversity of T cell memory.

In summary, our analyses of durable CD4^+^ T cell responses uncovered virus-specific CD4^+^ T cells that retain a naïve functional phenotype after vaccination. T_MN_ cells differ from T_SCM_ and other memory subsets by the lack of differentiation marker expression, yet they are antigen-experienced by TCR lineage analyses. The T_MN_ subset displays superior stability over time and is linked to durable and diverse T cell memory after vaccination. Understanding the generation, maintenance, and protective potential of T_MN_ cells could aid the future development of improved vaccine strategies for a broad range of pathogens.

### Limitation of Study

Our memory and naïve subsets are defined using phenotypic markers, without having examined their transcriptional or epigenetic states. While we have ruled out non-productive TCR engagement as a cause, how naïve phenotype is retained within a responding population remains unknown. Future studies will be needed to determine the differentiation trajectory toward T_MN_ state and if similar signals that drive T_SCM_ differentiation also promote T_MN_ development. As our analyses are focused on CD4^+^ T cell responses to YFV in healthy individuals, broader studies on CD8^+^ and CD4^+^ T cell responses to other pathogens would be needed to understand the prevalence of antigen-experienced T_MN_ cells and how they change with advanced age and disease. Notably, our data supporting T_MN_ in long-lived responses are correlational due to the nature of observational studies. Future investigations will be necessary to establish if T_MN_ cells directly contribute to durable immunologic memory, generate protective responses upon recall, and how they might be targeted to enhance the longevity of protective memory.

## Material and Methods

### Study Design

This study uses cryopreserved cells stored in fetal bovine serum (FBS) with 10% DMSO from an ongoing vaccine study at the University of Pennsylvania (*20*). This study includes 7 healthy adult participants with no prior YFV exposure who received one or two doses of the 17D live-attenuated YFV vaccine (YF-VAX®, Sanofi Pasteur). Five participants were followed longitudinally for 2 to 6.7 years after vaccination. All samples were de-identified and obtained with IRB regulatory approval from the University of Pennsylvania. Subject characteristics are shown in Table S1.

### Cell lines

Hi5 cells (ThermoFisher) were maintained by insect cell culture medium (ESF921, Expression Systems) supplemented with 0.02% gentamicin at 28℃.

### Protein expression and tetramer production

HIS-tagged HLA-DRA/B1*0301, 0401, 0407, and 1501 protein monomers of wild type (wt) sequence or with L112W, S118H, V143M, T157I mutations (*23*) were produced by Hi5 insect cells and extracted from culture supernatant using Ni-NTA (Qiagen). HLA-DR monomers were biotinylated overnight at 4°C using BirA biotin ligase (Avidity) and purified by size exclusion chromatography using Superdex 200 size exclusion column (AKTA, GE Healthcare). Biotinylation was confirmed by gel-shift assay. Peptide exchange and tetramerization for wildtype and modified affinity-matured DR were performed using standard protocols as previously described (*38, 39*). In brief, HLA-DR proteins were incubated with thrombin (Millipore) at room temperature for 3 - 4 hours and exchanged with peptides of interest in 50-fold excess at 37°C for 16 hours. Peptide-loaded HLA-DR monomers were incubated with fluorochrome-conjugated streptavidin at 4 - 5: 1 ratio for 2 min at room temperature, followed by a 15 min incubation with an equal volume of biotin-agarose slurry (Millipore). Tetramers were buffered exchanged into PBS, concentrated using Amicon ULTRA 0.5ml 100KDa (Millipore), and kept at 4 °C for no more than 2 weeks prior to use.

### *Ex vivo* T cell analyses and cell sorting

#### Phenotypic analyses and frequency quantification

Tetramer staining was performed on at least 10 million PBMCs with 5 ug of tetramers in 100 μl reaction for 1 hour at room temperature as previously described (*20, 39, 40*). Tetramer-tagged cells were enriched by adding anti-fluorochrome and anti-HIS MicroBeads (MiltenyiBiotec). The mixture was passed through LS columns (MiltenyiBiotec). Column-bound cells were washed and eluted according to manufacturer protocol. For antibody staining, the enriched samples were stained with viability dyes, exclusion markers (anti-CD19 and anti-CD11b, BioLegend), and surface markers (anti-CD3, anti-CD4, anti-CD45RO, anti-CCR7, anti-CD11a, anti-CD95, anti-CD49d, anti-CXCR3) in 50 to 100ul of FACS buffer (PBS plus 2% FCS, 2.5mM EDTA, 0.025% Sodium Azide) for 30 minutes at 4°C. Samples were fixed with 2% paraformaldehyde and acquired by flow cytometry using LSRII (BD). Data analyses were performed by FlowJo (BD). Frequency calculation was obtained by mixing 1/10th of samples with 200,000 fluorescent beads (Spherotech) for normalization.

For longitudinal experiments involving both wt and modified DR, paired data from wt and mutant DR, with a minimum of two data points per time point for each specificity, were used to derive the equation for normalization: log_2_(Freq_modified_)=3.72+0.35 * log_2_(Freq_wt_) (Fig. S4D). Frequencies generated by wt tetramers that were below the normalized values were adjusted. Mixed effects exponential decay models were used to analyze longitudinal changes in antigen-specific T cell populations and estimate the corresponding slopes. These models were implemented in *MonolixSuite 2021R1* (Lixoft) and fitted to data after vaccination. Initial T cell specificity values were lognormally distributed, exponential decay rates were normally distributed, and lognormal multiplicative error was used. The estimation of the population parameters was performed using the Stochastic Approximation Expectation-Maximization (SAEM) algorithm. Half-lives were calculated as ln(2)/*k*, where the corresponding *k* values represented the estimated exponential decay rate constants. Estimated decay rates were converted into slopes as -*k*.

For multi-dimensional analyses, a total of 1465 manually gated tetramer^+^ cells were exported from FlowJo, read into R by flowCore, and combined into one single dataset for subsequent data processing and analyses using the Spectre package in R (*21*). Staining intensities were converted using Arcsinh transformation with a cofactor of 200. Batch alignment was performed using the CytoNorm (*41*). Clustering was performed using Phenograph with nearest neighbors set to 55 (*k* = 55) (*42*). UMAP was used for dimensional reduction and visualization (*43*).

#### Function response

T cells were rested overnight, followed by 4 - 5 hours of stimulation by phorbol myristate acetate (PMA, 5 ng/ml, Sigma) and ionomycin (500 ng/ml, Sigma) in the presence of monensin (2 uM, Sigma) and Brefeldin A (5 ug/ml, Sigma). Tetramer and surface antibody staining were performed as above.

Intracellular staining with antibodies to TNF-α, IFN-γ, IL-2, CD3 and CD4 (Biolegend) was performed following BD Cytofix/Cytoperm Fixation/Permeabilization Kit according to manufacturer protocol (BD).

#### Cell sorting

Cell numbers were increased to around 60 million CD3^+^ or CD4^+^ T cells, stained in up to 10 ug of each tetramer in a 100 ul reaction. Antibody staining was performed as above without fixation. Individual tetramer-labeled cells were isolated for TCR sequencing or T cell cloning by index sorting using the purity mode on FACS Aria (BD).

### Generation and stimulation of T cell clones

#### Clone generation

Cells were stained with tetramers and enriched with magnetic beads as described above. Single tetramer-stained CD4^+^ T cells were sorted into individual wells in a round bottom 96-well plate containing 10^5^ irradiated PBMCs, 10^4^ JY cell line (ThermoFisher), PHA (1:100, ThermoFisher), IL-7 (25 ng/ml, PeproTech), and IL-15 (25 ng/ml, PeproTech). IL-2 (50 IU/ml, PeproTech) was added on day 5 and replenished every 3-5 days. Cells were resupplied with fresh medium with IL-2 (50 IU/ml), PHA (1:100), and 10^5^ irradiated PBMCs every two weeks.

#### DCs generation

Monocytes from HLA-DR allele-matched donors were isolated using negative enrichment kits (RosetteSep Human Monocyte Enrichment Cocktail, StemCell). 10 million cryopreserved monocytes were cultured in 15 ml DC media (RPMI 1640 plus Glutamine,10% FCS, 1X Pen/Strep, 10 mM HEPES) in the presence of 100 ng/ml GM-CSF and 500 U/ml IL-4. Three days later, half the culture media was replaced with fresh DC media with 100 ng/ml GM-CSF, 500 U/ml IL-4, and 0.05 mM 2-mercaptoethanol. Cells in suspension were harvested at 5 to 6 days and added to a flat-bottom 96-well plate at 25,000 DCs per well. DCs were treated with 100 ng LPS and peptides (0.00001 ug/ml to 10ug/ml) for 16 hours and replenished with fresh media before co-culturing with T cells.

#### Stimulation of T cell clones

T cell clones were rested overnight in fresh media without IL-2 and added to wells containing matured DCs at 1:1 ratio in the presence of monensin (2 uM, Sigma) and Brefeldin A (5 ug/ml, Sigma). After 5 hours, cells were transferred into a new 96-well round bottom plate, washed once with FACS buffer, and stained with viability dyes, exclusion markers (anti-CD19 and anti-CD11b, BioLegend) for 30 minutes at 4°C. Intracellular staining with antibodies to TNF-α, IFN-γ, IL-2, CD3 and CD4 (Biolegend) was performed following BD Cytofix/Cytoperm Fixation/Permeabilization Kit according to manufacturer protocol (BD). Half maximal effective concentration (EC50) was determined using the percentage of T cell clones that produced TNF-α in response to decreasing peptide concentrations (10, 1, 0.1, 0.01, 0.001, 0.0001, and 0.00001 ug/ml). A non-linear fit without constraint was applied to log-transformed concentration using the equation Y=Bottom + (Top-Bottom)/(1+10^((LogEC50-X)*HillSlope)) in Prism (GraphPad). For proliferation assay, T cell clones were labeled with 1:1000 diluted CellTrace Violet (CTV, ThermoFisher Scientific) following manufacturer protocol. The CTV-stained cells were rested in fresh media without IL-2 for 16 hours. 25,000 rested T cells were co-cultured with DC pulsed with 10 ug/ml cognate peptides or treated with PHA as a positive control (1:100, ThermoFisher). After 5 days, cells were harvested and stained with viability dyes and surface antibodies (anti-CD19, anti-CD11b, anti-CD3, and anti-CD4, BioLegend) for 30 minutes at 4°C followed by fixation with 2% paraformaldehyde. Samples were acquired by flow cytometry using LSRII (BD) and analyzed by FlowJo (BD).

### Single-cell TCR sequencing and analyses

Single-cell TCR Sequencing by nested PCRs was performed using the primer sets and the protocol as previously described (*20, 44*). In brief, reverse transcription was performed with CellsDirect One-Step qRT-PCR kit according to the manufacturer’s instructions (CellsDirect, Invitrogen) using a pool of 5’ TRVB-region specific primers and 3’ C-region primers. The cDNA library was amplified using a second set of multiple internally nested V-region and C-region primers with HotStarTaq DNA polymerase kit (Qiagen). The final PCR reaction was performed on an aliquot of the second reaction using a primer containing common base sequence and a third internally nested Cβ primer. PCR products were gel purified (Qiagen) and sequenced on Novaseq 6000 platform (Illumina). TCR sequences were pre-processed as previously described (*20*). In brief, forward and reverse reads were converted into one paired end read using pandaseq (*45*). Data were demultiplexed by the unique combination of plate, row, and column barcodes. Consensus TCRβ sequences were identified using the V(D)J alignment software MiXCR (*46*). A threshold of a read count of 200 reads per sequence was applied to the consensus sequences. If more than one TCRa or TCRý chain passes this criterion we retain the dominant TCRβ and the two TCRa chains with the highest read count. For data obtained from cells several months after vaccination, we additionally require phenotypic annotation based on antibody staining from index sort data. Data were excluded if phenotypic information was not retained or ambiguous. For downstream analyses, data wrangling was performed using the tidyverse package. TCRs were matched by TCRβ if only the beta chain was available, or by TCRβ plus at least one TCRα if alpha chain(s) were called. Circos plots were made using the circlize package of R software (*47*).

### Statistical Methods

Normality was assessed using D’Agostino-Pearson test. Spearman was used if either of the two variables being correlated was non-normal. Otherwise, Pearson was used to measure the degree of association. Least squares linear regression was used to calculate the best-fitting line. Statistical comparisons were performed using two-tailed Student’s t-test, paired t-test, Welch’s one-way ANOVA, repeated measures one-way ANOVA, two-way ANOVA, or mixed effects model. A p-values of <0.05 was used as the significance level and adjusted if multiple comparisons were performed. Statistical analyses were performed using GraphPad Prism. Lines and bars represent the mean and variability is represented by the standard error of the mean (SEM). * P < 0.05, ** P < 0.01, *** P < 0.001, **** P < 0.0001.

## Supplementary Materials

**Figure S1:**
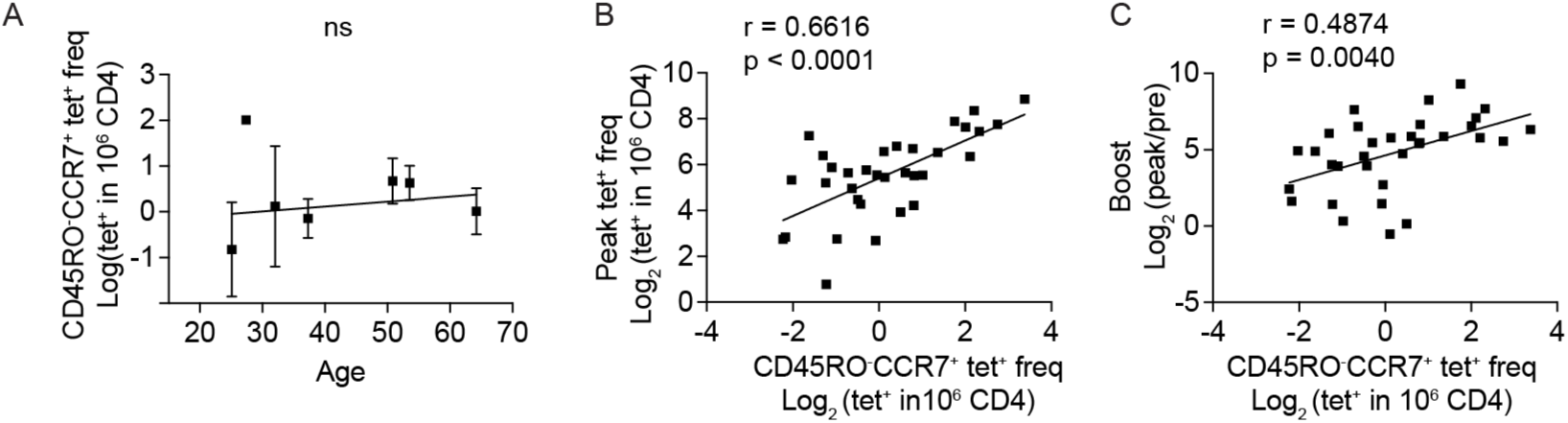
CD45RO^-^CCR7^+^ YFV-specific CD4^+^ T cells by age and their relationship to the effector response. (A) The frequency of post-immune CD45RO^-^CCR7^+^ YFV tetramer^+^ CD4^+^ T cells in relationship to donor age. Distinct tetramer^+^ populations from the same donor are combined and represented as an average (n = 7). (B) The correlation between the frequency of CD45RO^-^CCR7^+^ YFV tetramer^+^ cells measured at least 7 months after vaccination and the highest total tetramer^+^ frequency from a previous time point measured within the first month after vaccination (n = 36). (C) The correlation between CD45RO^-^CCR7^+^ YFV tetramer^+^ T cell frequency at least 7 months after vaccination and the fold-change between peak frequency and the pre-vaccine baseline. Pearson correlation was computed.

**Figure S2:**
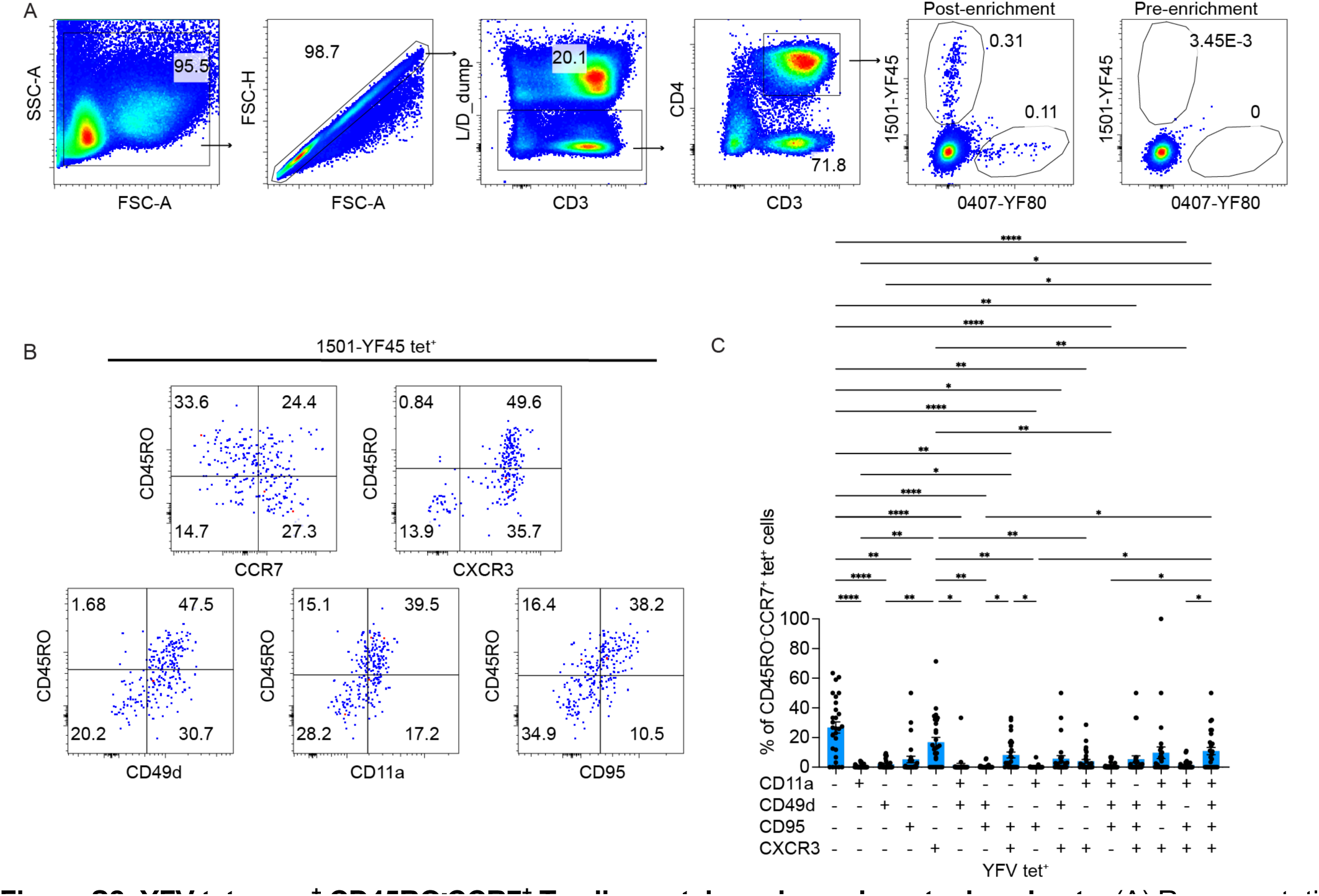
YFV tetramer^+^ CD45RO^-^CCR7^+^ T cells contain various phenotypic subsets. (A) Representative plots show the gating strategy used to identify tetramer^+^ cells. (B) FACS plots show the phenotype of YF45-specific T cells by the indicated antibody staining. (C) Boolean gates for CD11a, CD49d, CD95, and CXCR3 were applied onto manually gated CD45RO^-^CCR7^+^ YFV tetramer^+^ T cells. The plot shows various phenotypic combinations. Each symbol represents a tetramer^+^ population (n = 28). One-way ANOVA was performed and corrected with Tukey’s multiple comparison test.

**Figure S3:**
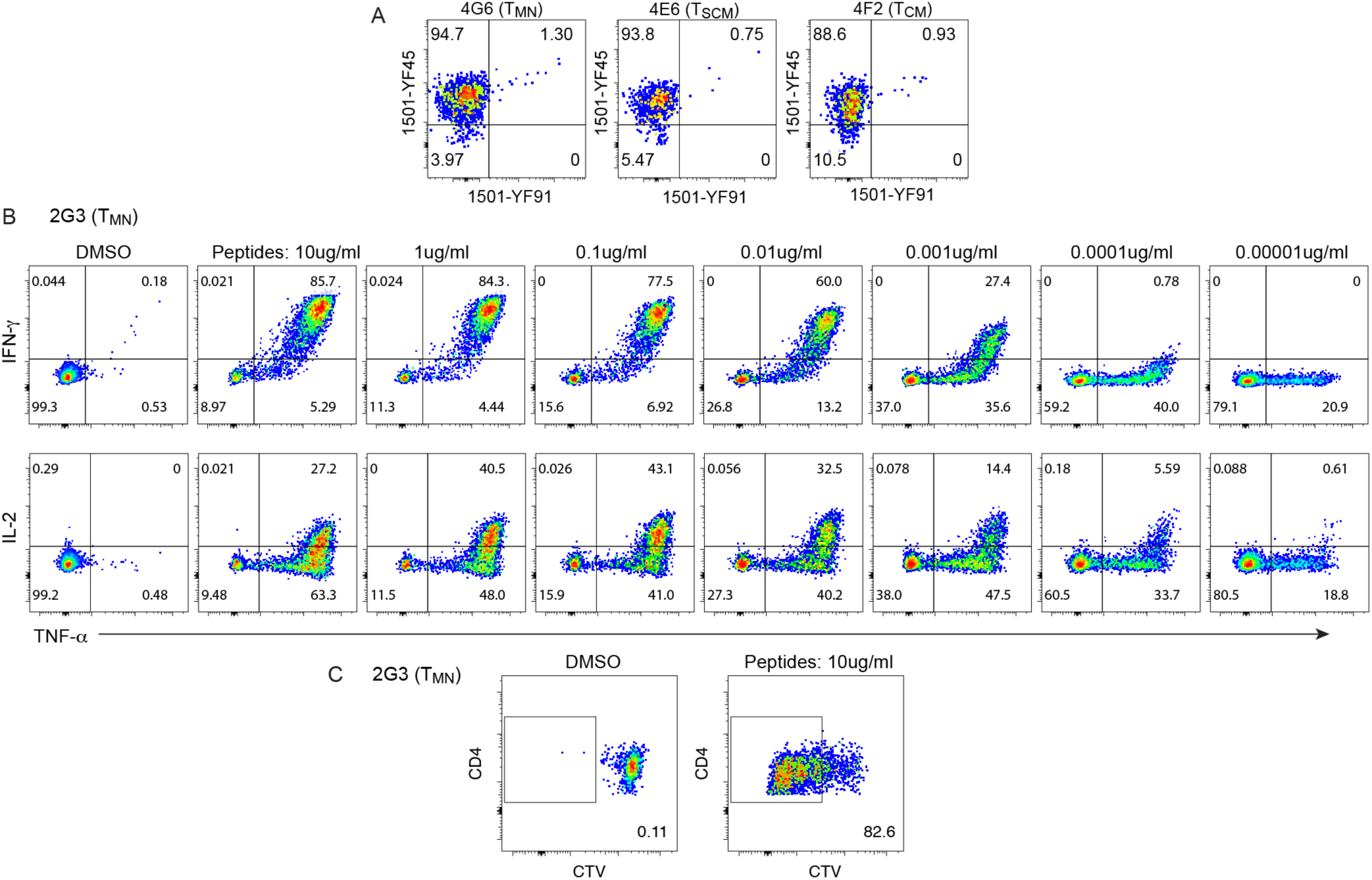
Tetramer staining and peptide responses of YFV-specific T cell clones. (A) Representative tetramer staining of T_MN_, T_SCM_, and T_CM_-derived YFV-specific T clones. (B) Plots show cytokine response by a T_MN_ clone to decreasing concentrations of the cognate YF45 peptide after a 5-hour co-culture with peptide-loaded DCs. (C) Representative plots show CTV staining of a CTV-labeled T cell clone after a 5-day culture with vehicle-treated or peptide-loaded DCs.

**Figure S4:**
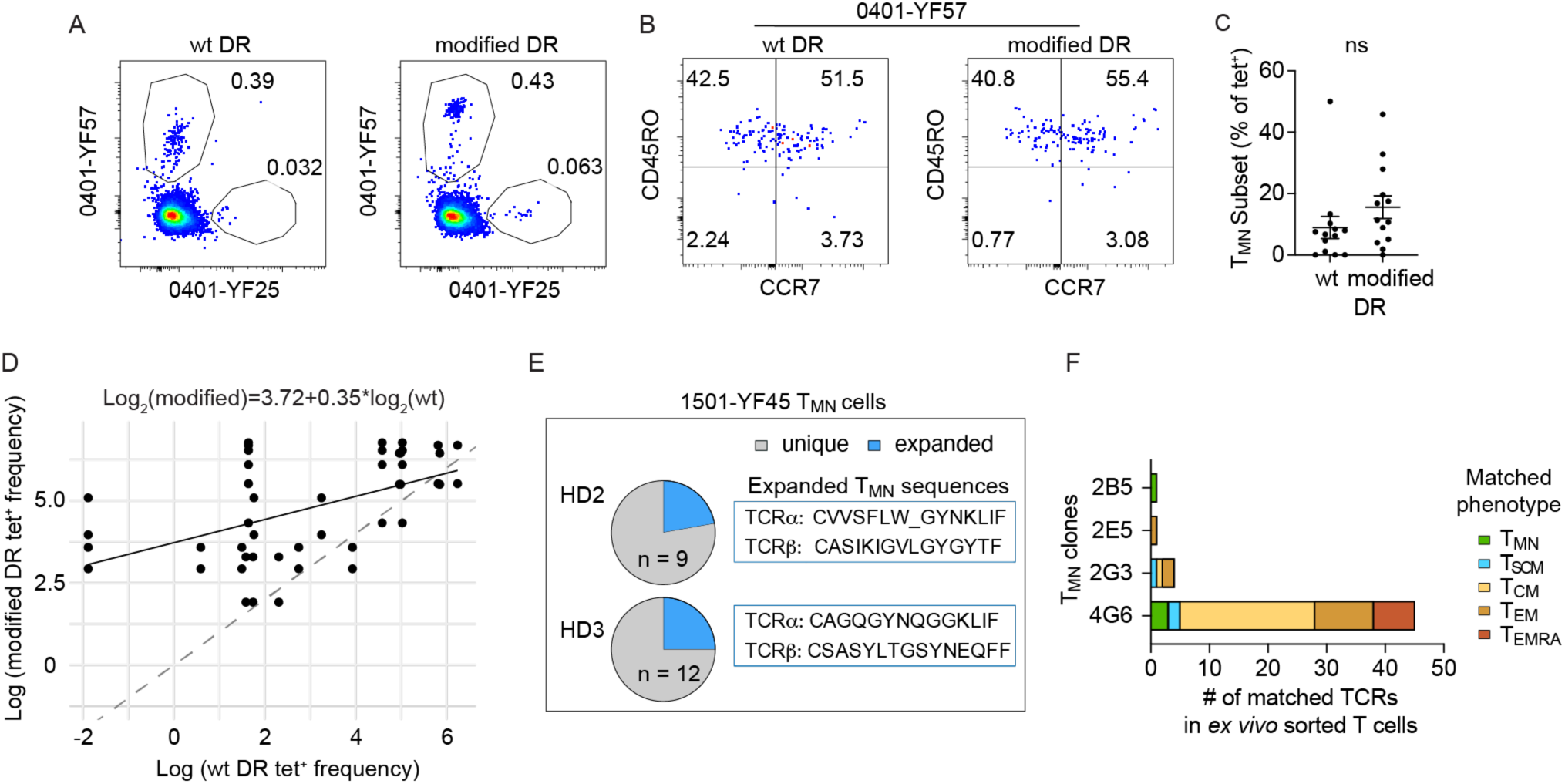
T _MN_ cells are antigen-experienced. (A) Representative plots show the identification of YFV-specific T cells using tetramers generated from wt or modified DR (L112W, S118H, V143M, T157I). (B) CD45RO and CCR7 expression of YF57-specific T cells stained by wt or modified DR. (C) Quantification of T_MN_ fraction within tetramer^+^ populations identified by wt or modified tetramers using the same sample. (D) The plot displays paired data from wt and modified DR, with a minimum of two data points per time point for each specificity, which were used to derive the equation for normalization (top). (E) Pie-charts show the distribution of unique versus expanded clonotypes within T_MN_ subset of YF45-specific T cells obtained 210-242 days after YFV vaccination. The sequences of the expanded clonotype from each donor are as indicated. (F) T_MN_-derived T cell clones were re-stained with tetramers and sorted for TCR sequencing. Clone-derived TCRs were compared with sequences from directly sorted T cells. The graph shows clones with matched TCRs and indicates the number of matched cells and their associated *ex vivo* phenotypes. For (C), Wilcoxon matched-pairs signed rank test was performed.

**Figure S5:**
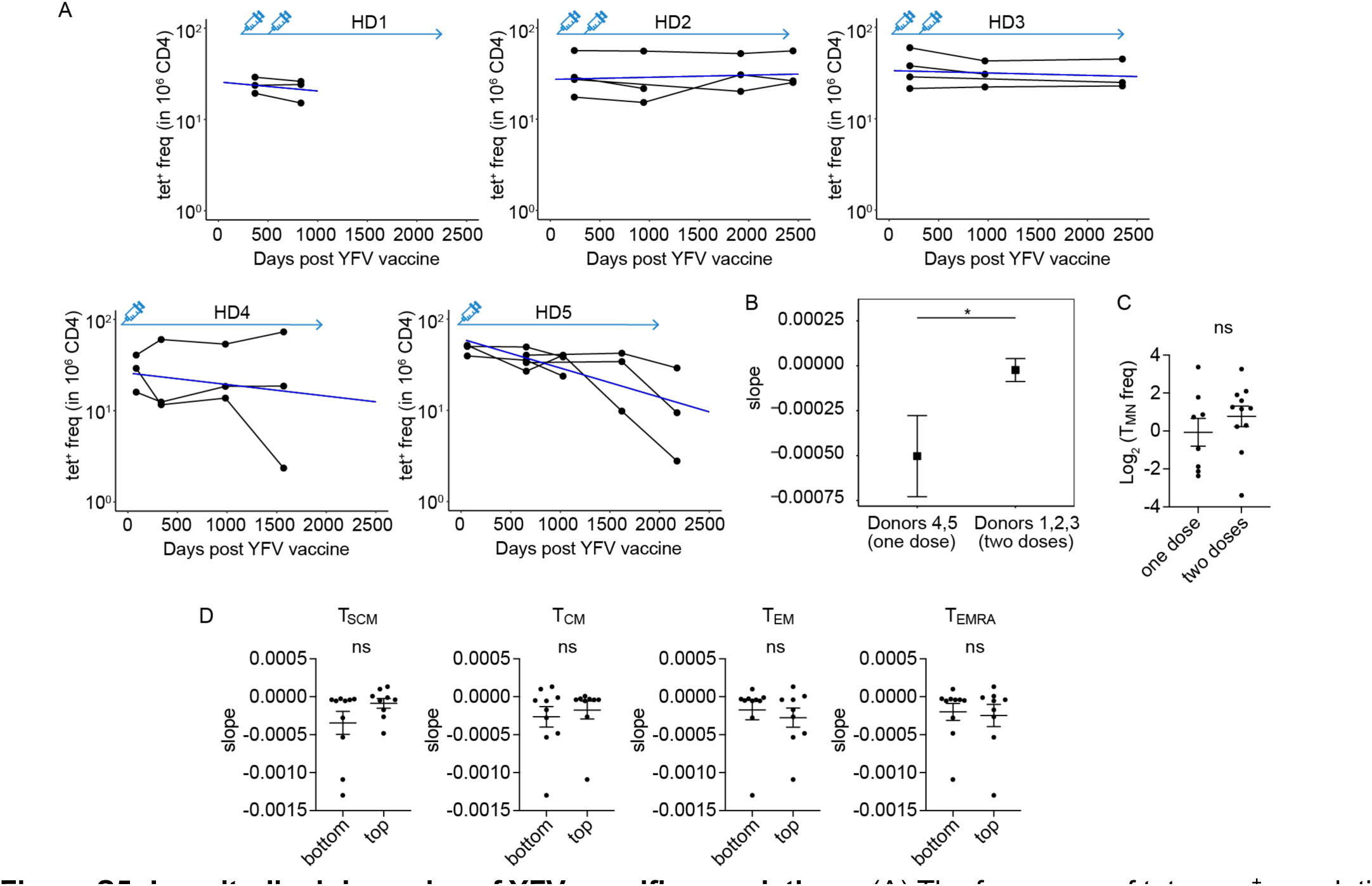
Longitudinal dynamics of YFV-specific populations. (A) The frequency of tetramer^+^ populations by donor. Filled circles represent individual tetramer^+^ populations. A line connects T cells labeled by the same tetramer across time points. Mixed effects exponential decay models were fitted to the longitudinal frequencies of individual tetramer-labeled populations for each donor. Blue lines represent the estimated decay. (B) Longitudinal dynamics of tetramer^+^ populations were combined for donors who received one YFV dose (donors 4 and 5) and two doses (donors 1, 2, 3). A mixed-effects exponential decay model was employed to estimate the corresponding population slopes for the two groups. (C) The plot summarizes T_MN_ frequency in donors who received one (donors 4, 5) or two doses (donors 1, 2, 3) of the YFV vaccine. (D) Plots summarize the estimated slopes of individual tetramer^+^ populations, divided into top and bottom halves by the averaged frequency of cells expressing the indicated phenotypes. (B) Wald test was used. (C) Mann-Whitney test was used. (D) Welch’s t-test was used.

**Table S1:** Donor characteristics

| ID | Sex | Age at the time of YFV vaccination | Vaccine dates | Sample collection dates |
| --- | --- | --- | --- | --- |
| HD1 | M | 37 | 8/25/16 | 8/28/17 |
| HD2 | F | 32 | 9/12/16 | 5/12/17 |
| HD3 | M | 64 | 1/4/17 | 8/2/17 |
| HD4 | M | 25 | 5/22/17 | 4/23/18 |
| HD5 | F | 50 | 4/21/17 | 2/7/19 |
| HD6 | M | 53 | 3/6/17 | 10/24/19 |
| HD7 | M | 27 | 3/6/17 | 2/23/21 |

**Table S2:**
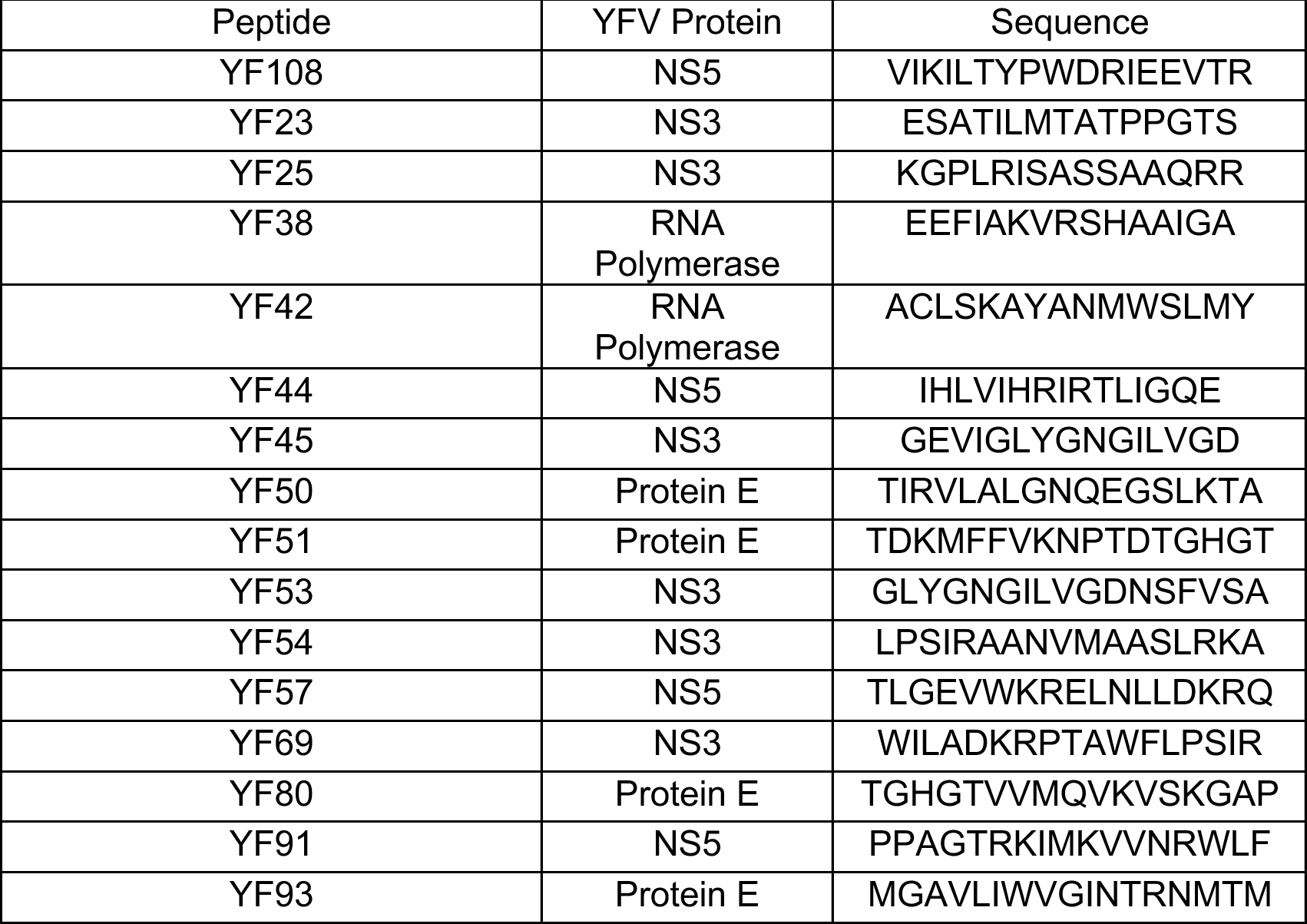
List of YFV peptides

**Table S3:** phenotypic markers

| Phenotype | CD45RO and CCR7 staining | CD95, CXCR3, CD49d, CD11a staining |
| --- | --- | --- |
| Naive | CD45RO-CCR7+ | not available |
| T<sub>CM</sub> | CD45RO+CCR7+ |  |
| T<sub>EM</sub> | CD45RO+CCR7- |  |
| T<sub>EMRA</sub> | CD45RO-CCR7- |  |
| T<sub>SCM</sub> | CD45RO-CCR7+ | positive for one of the following: CD95, CXCR3, CD49d, CD11a |
| T<sub>MN</sub> | CD45RO-CCR7+ | negative for all of the following: CD95, CXCR3, CD49d, CD11a |

**Table S4:**
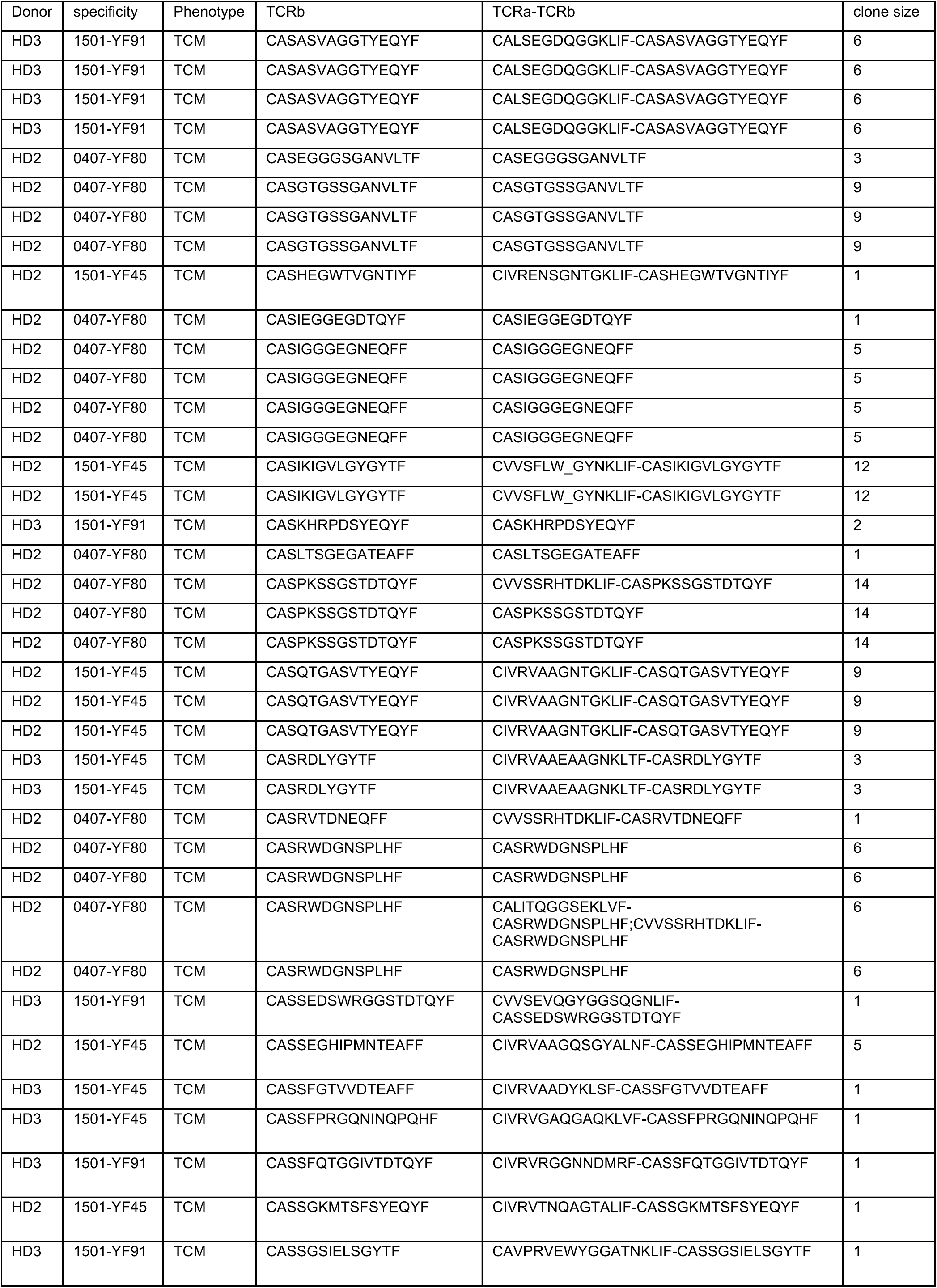
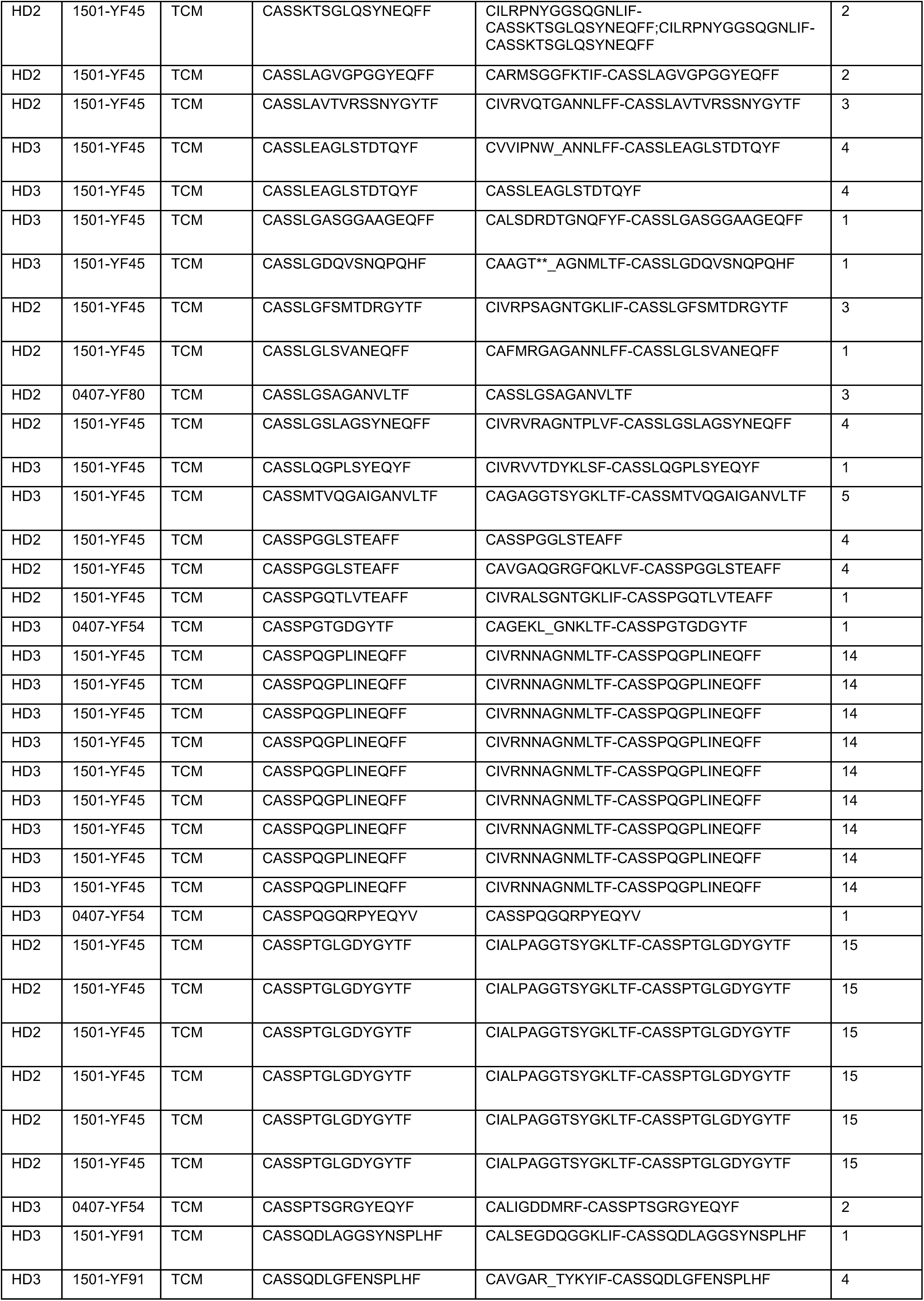
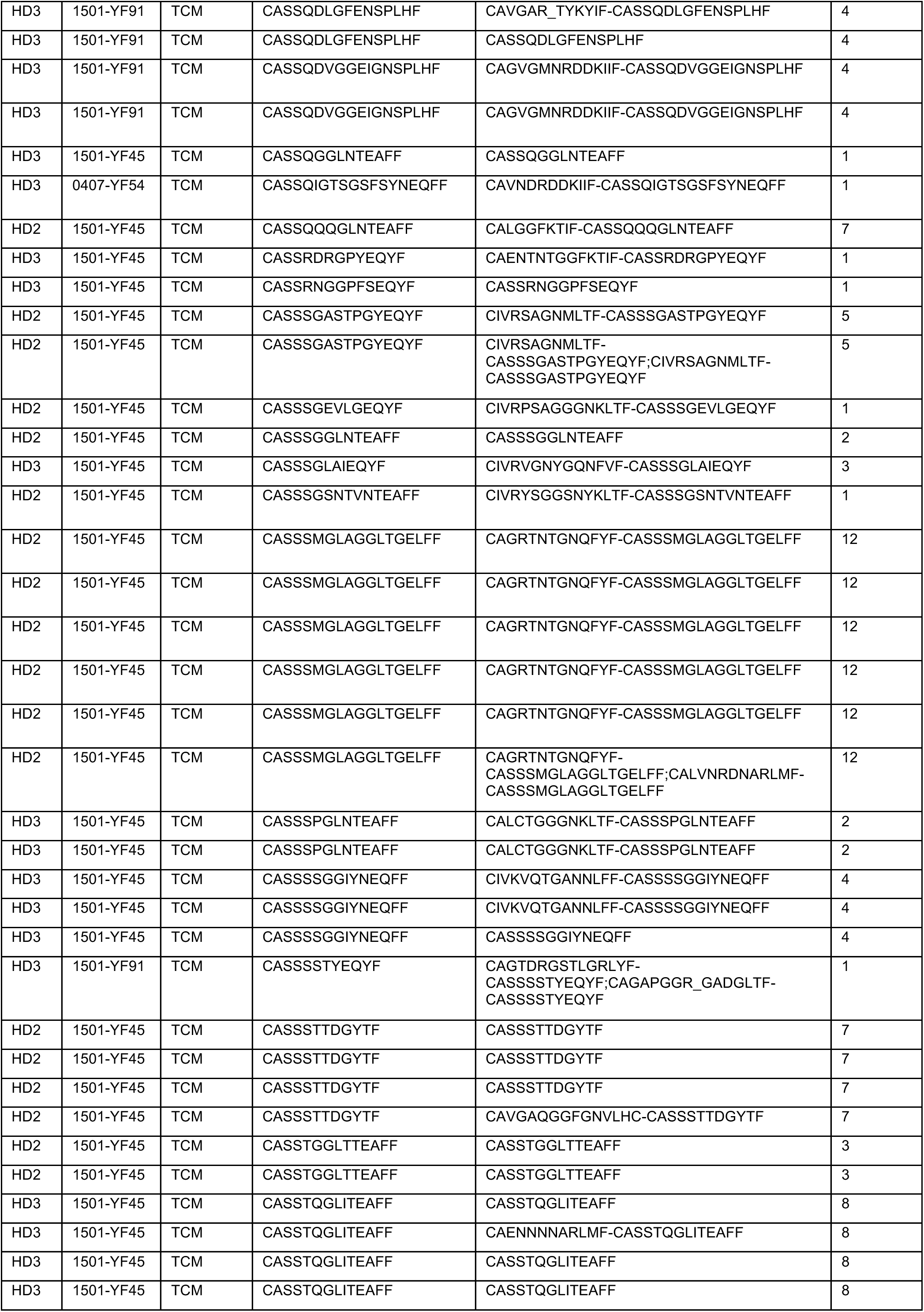
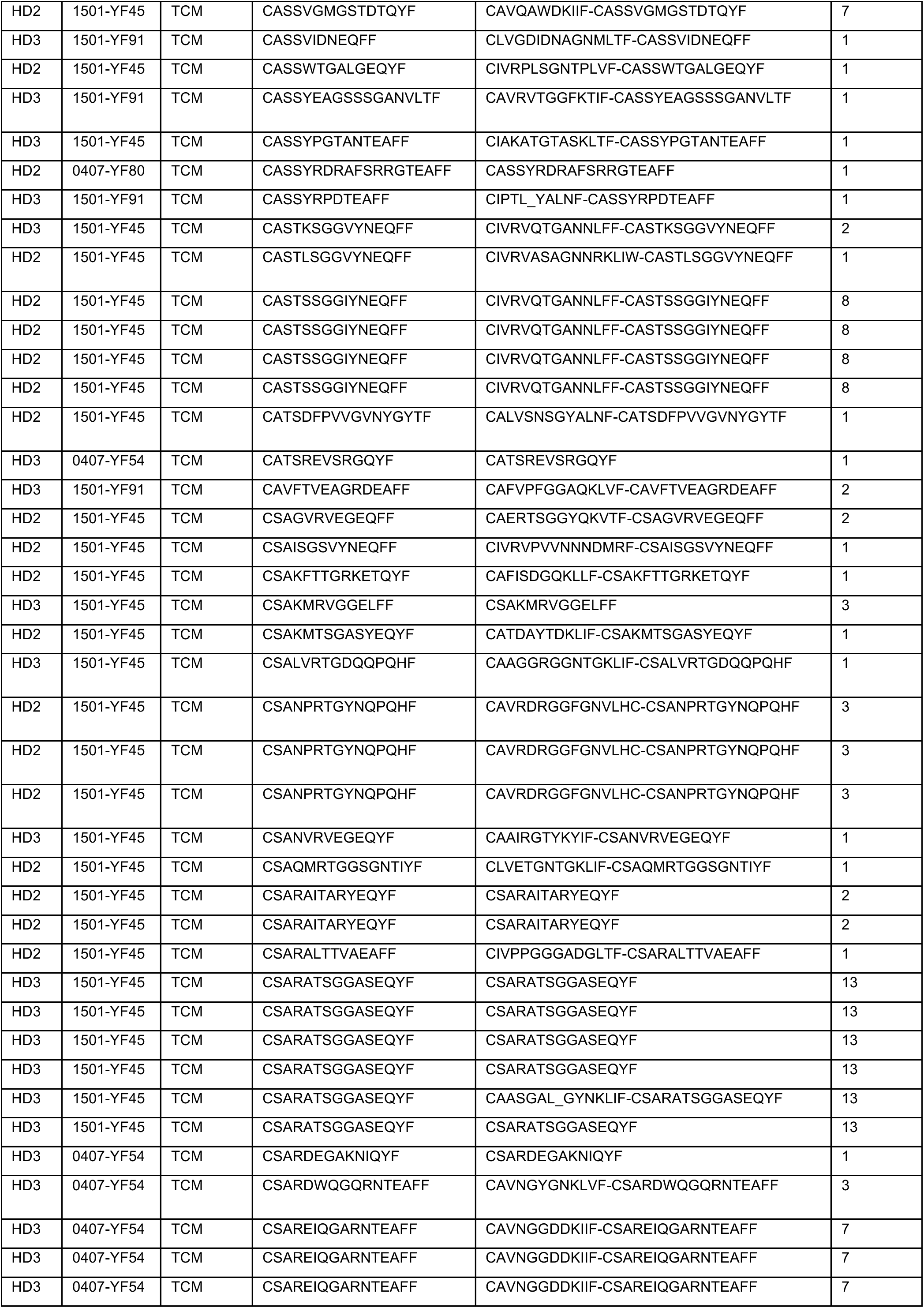
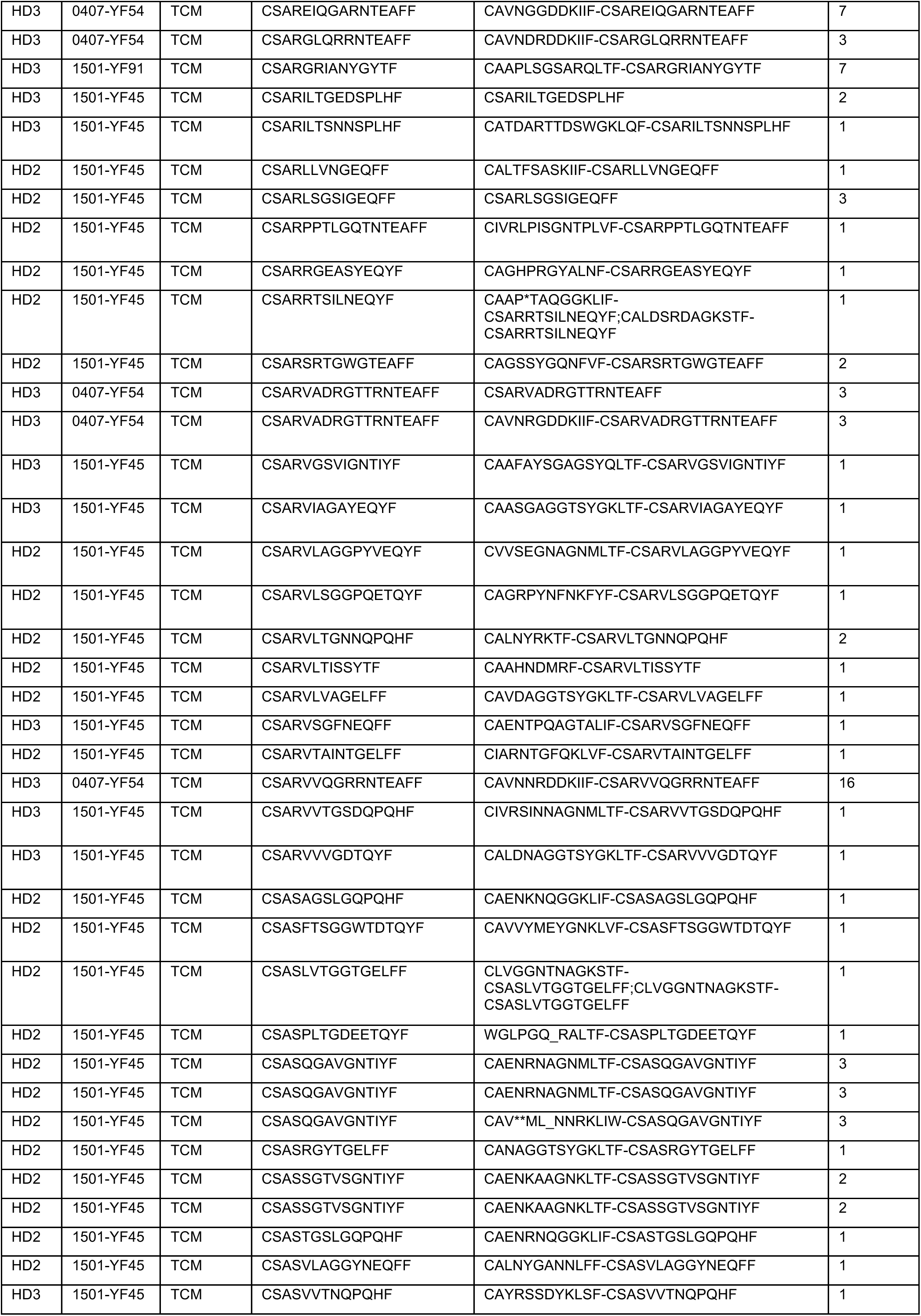
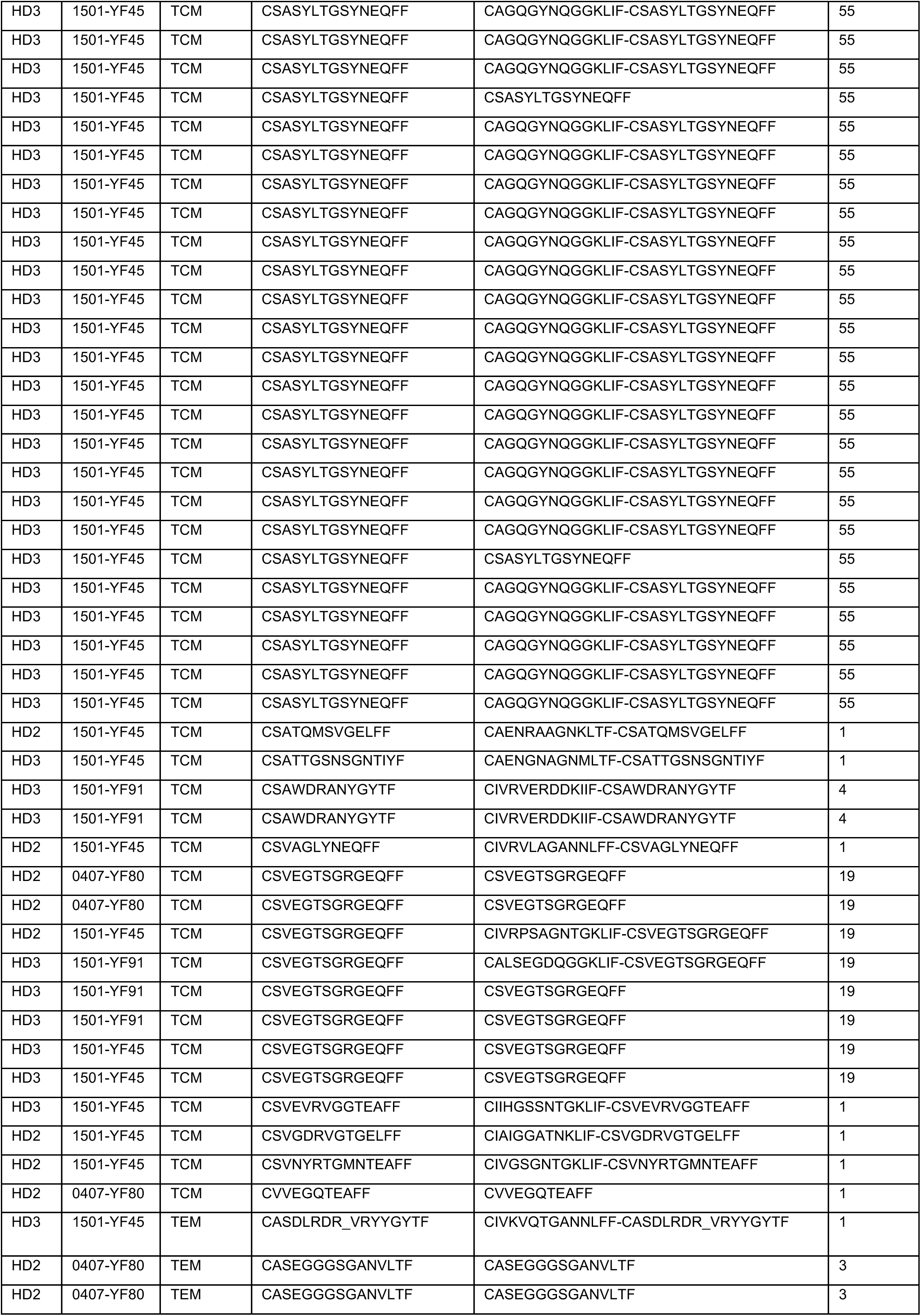
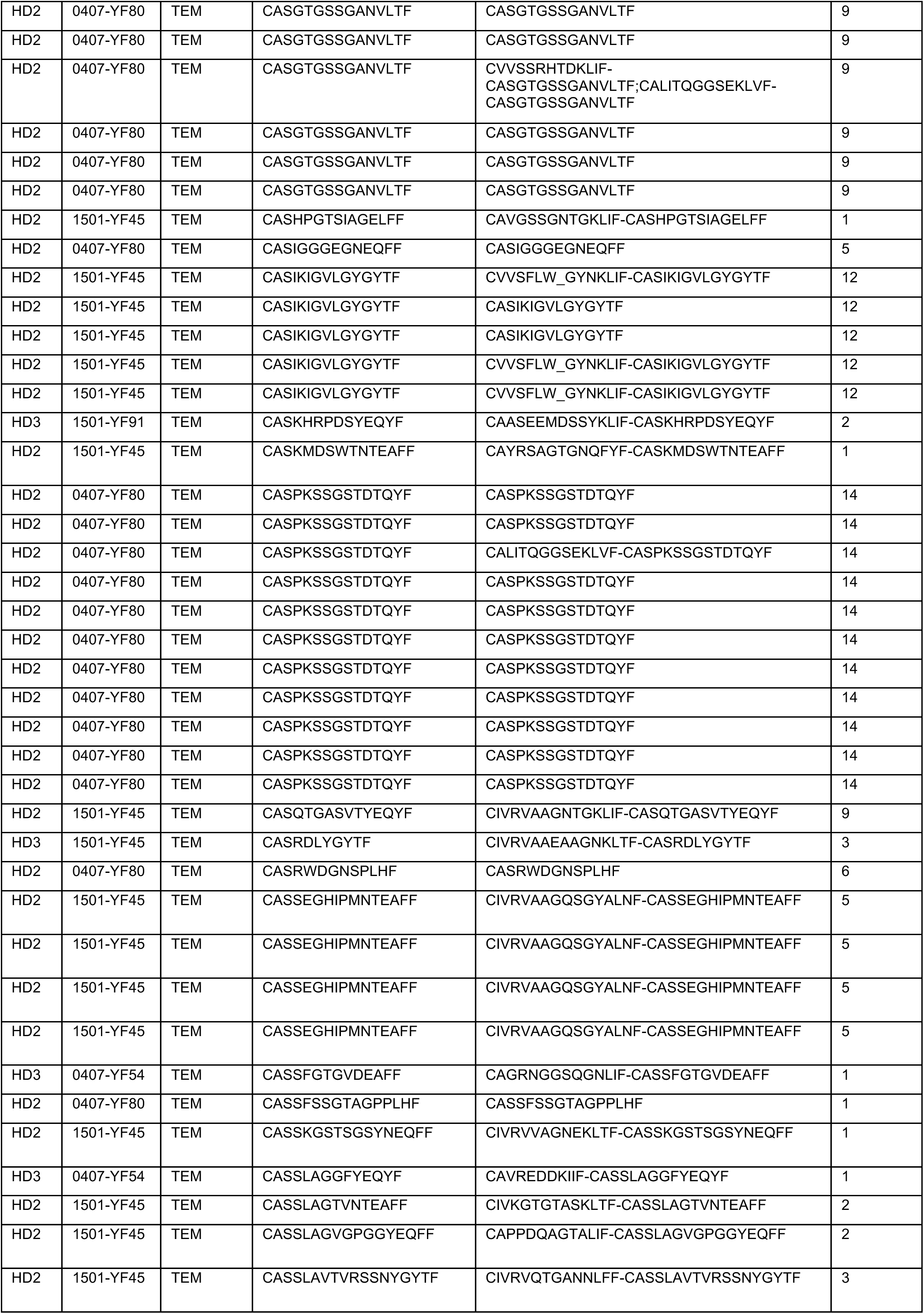
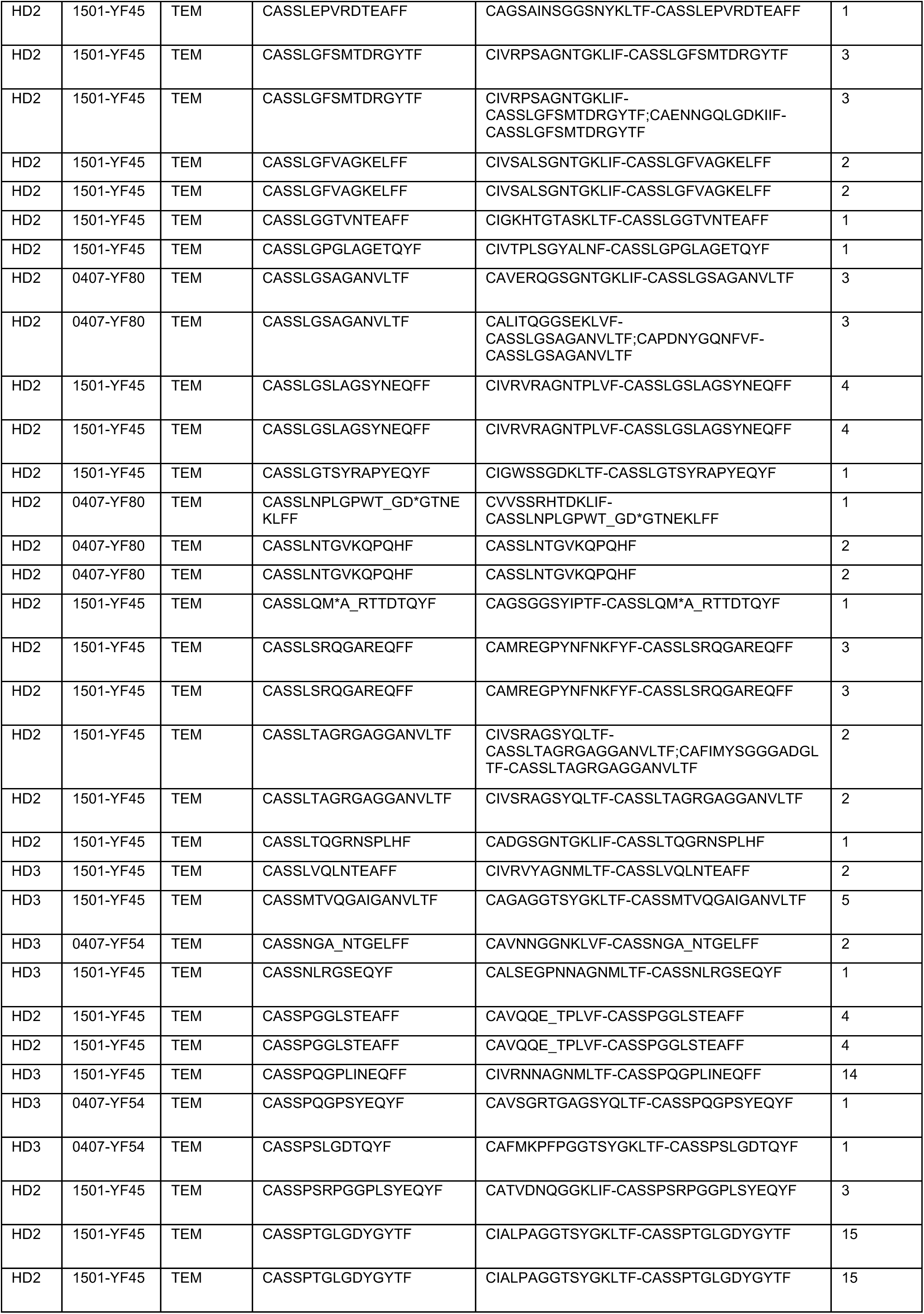
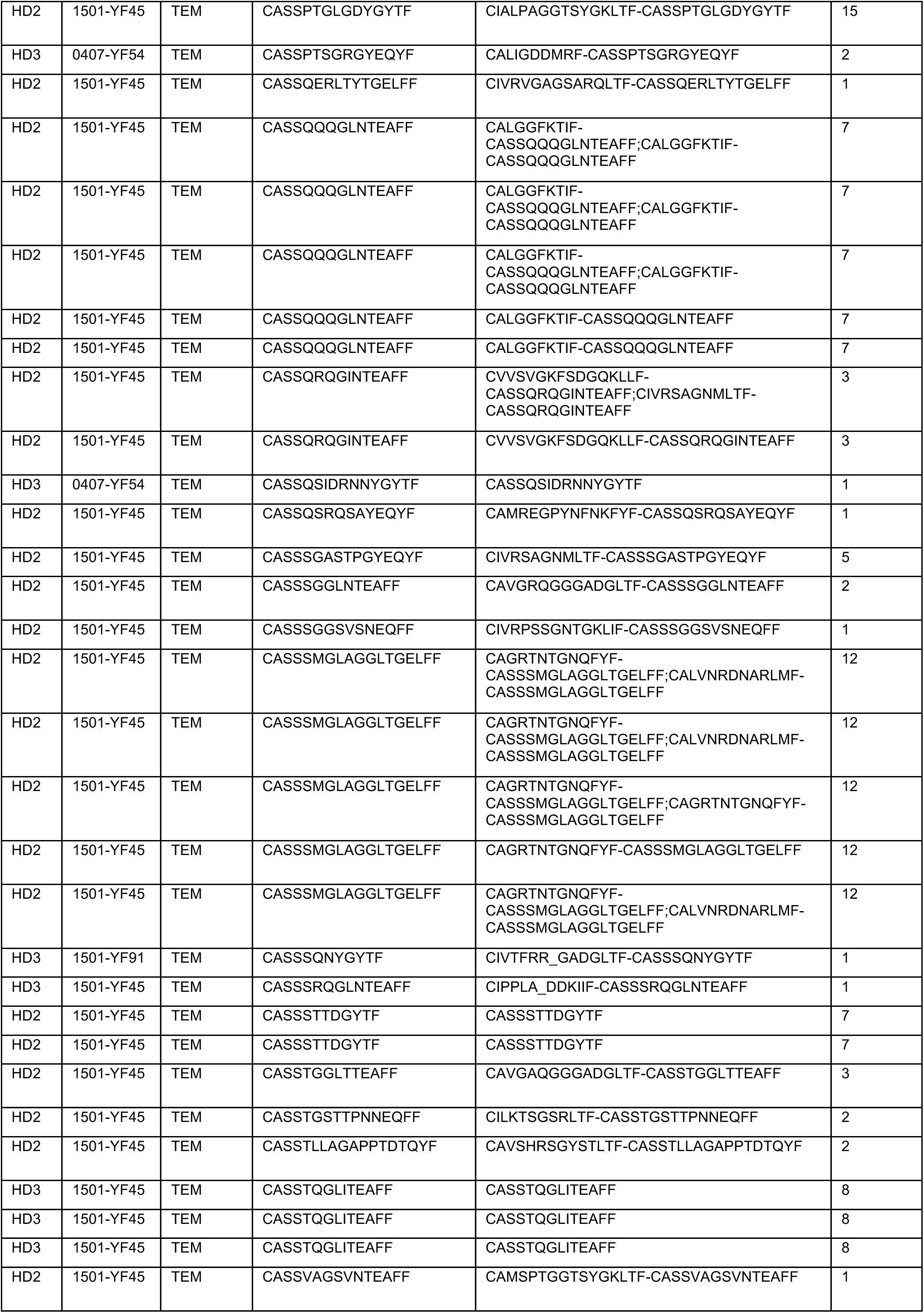
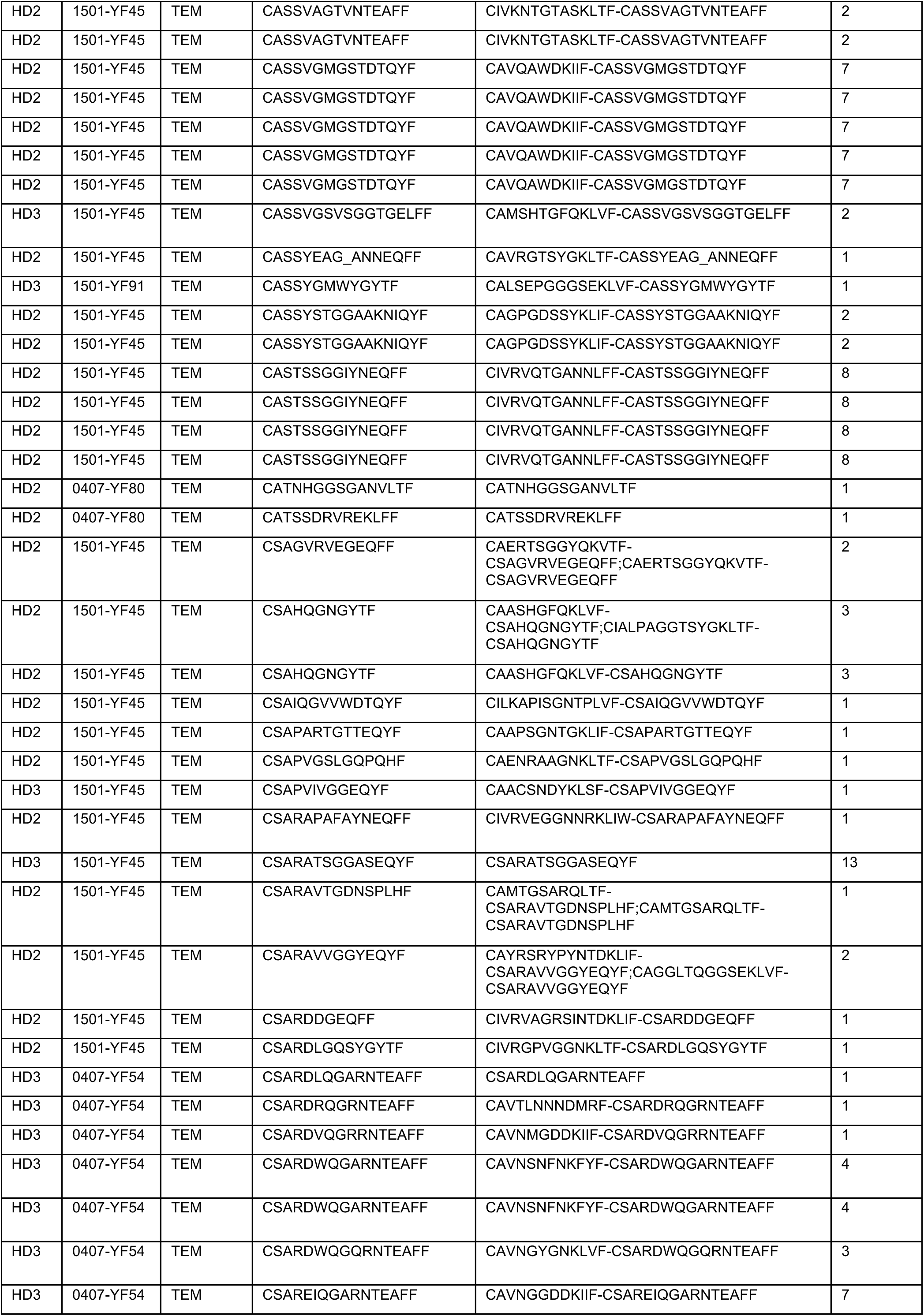
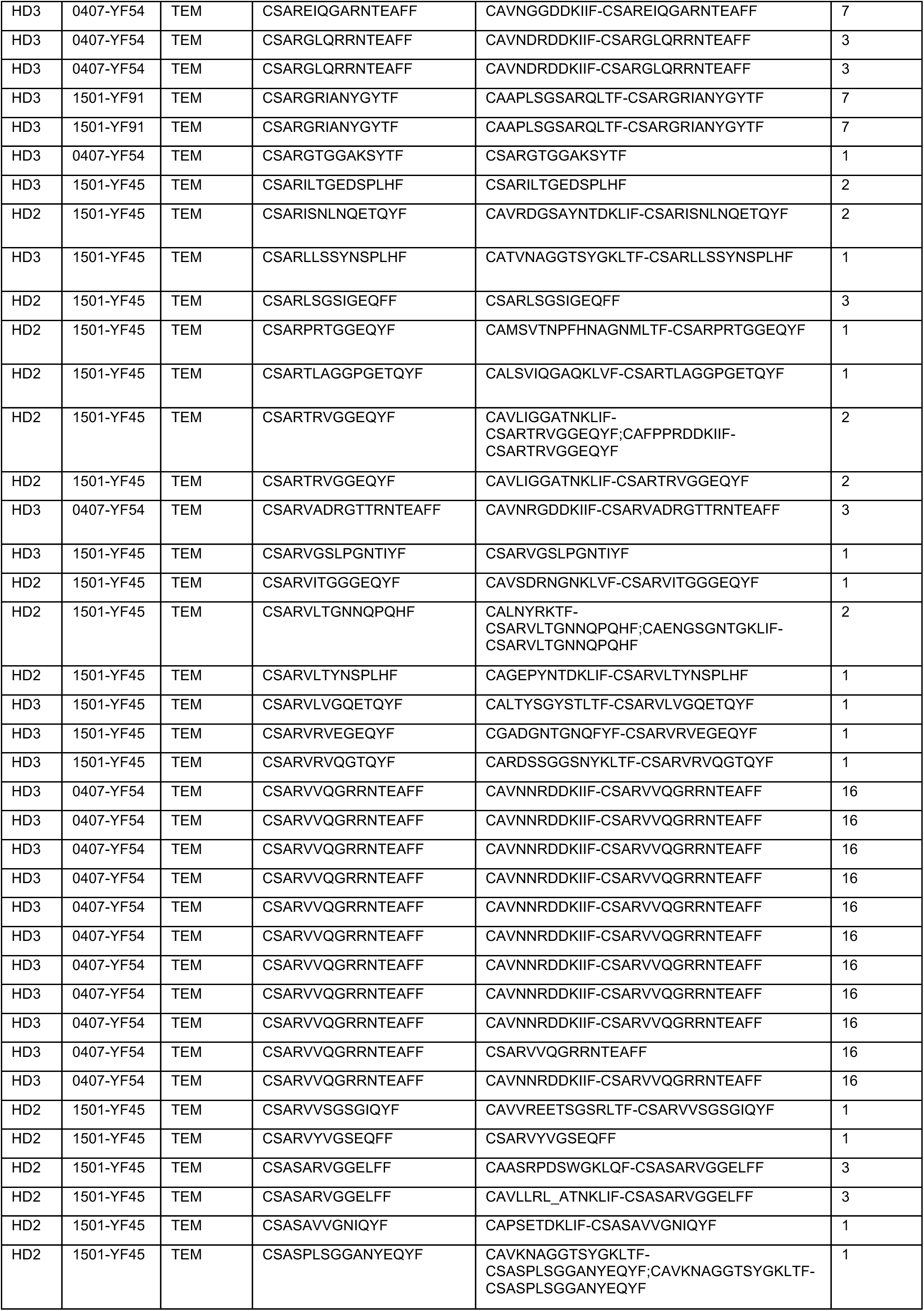
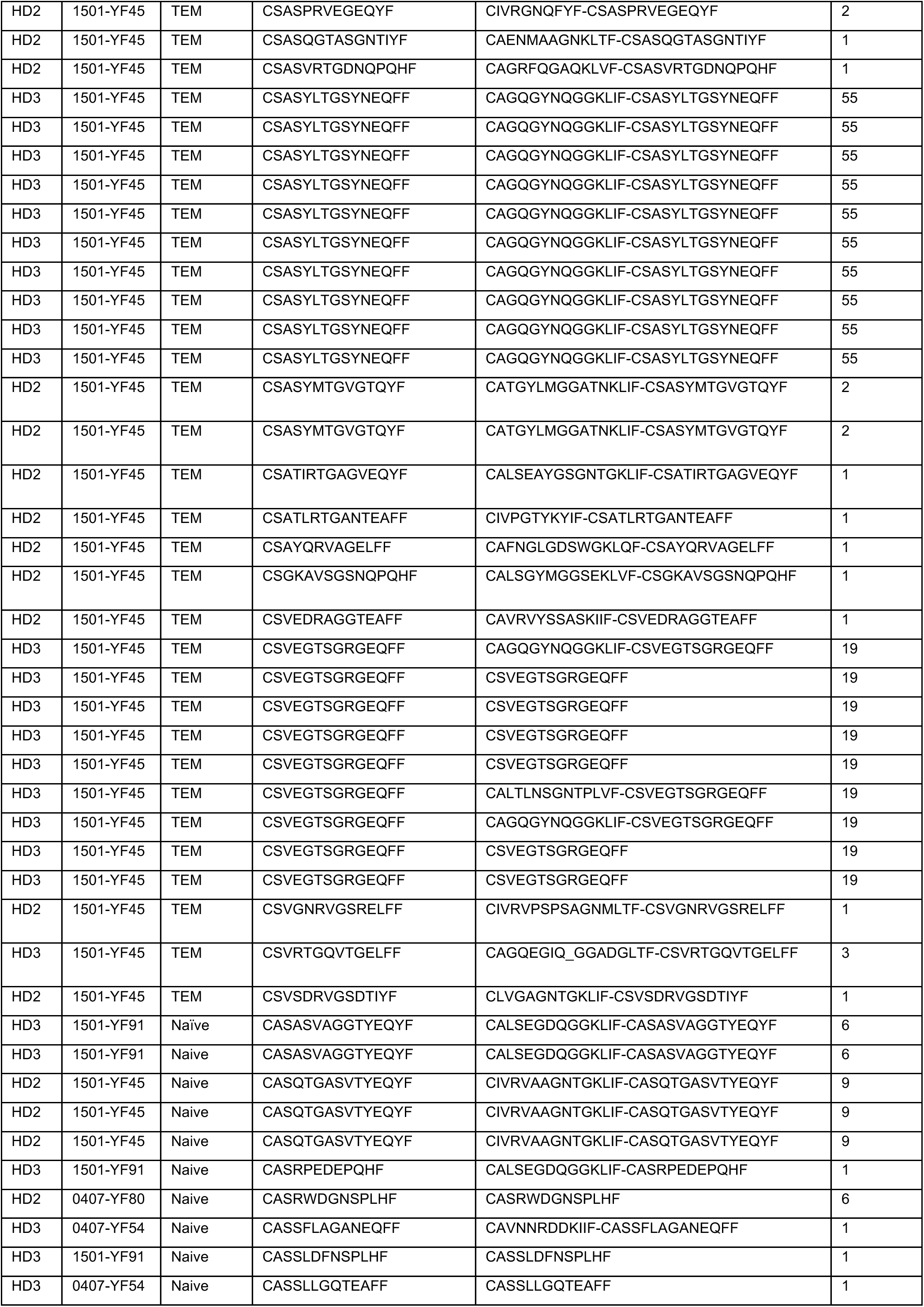
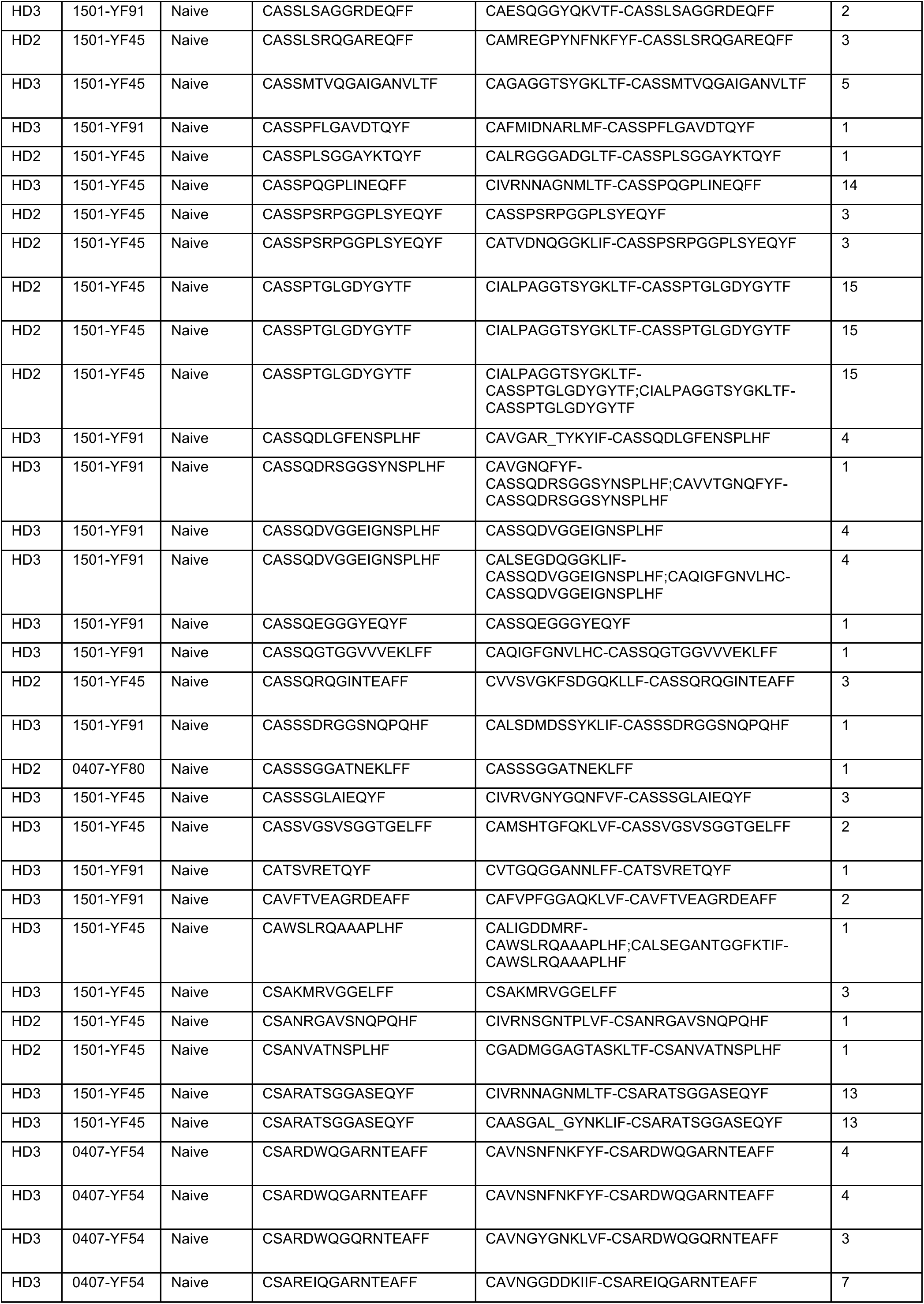
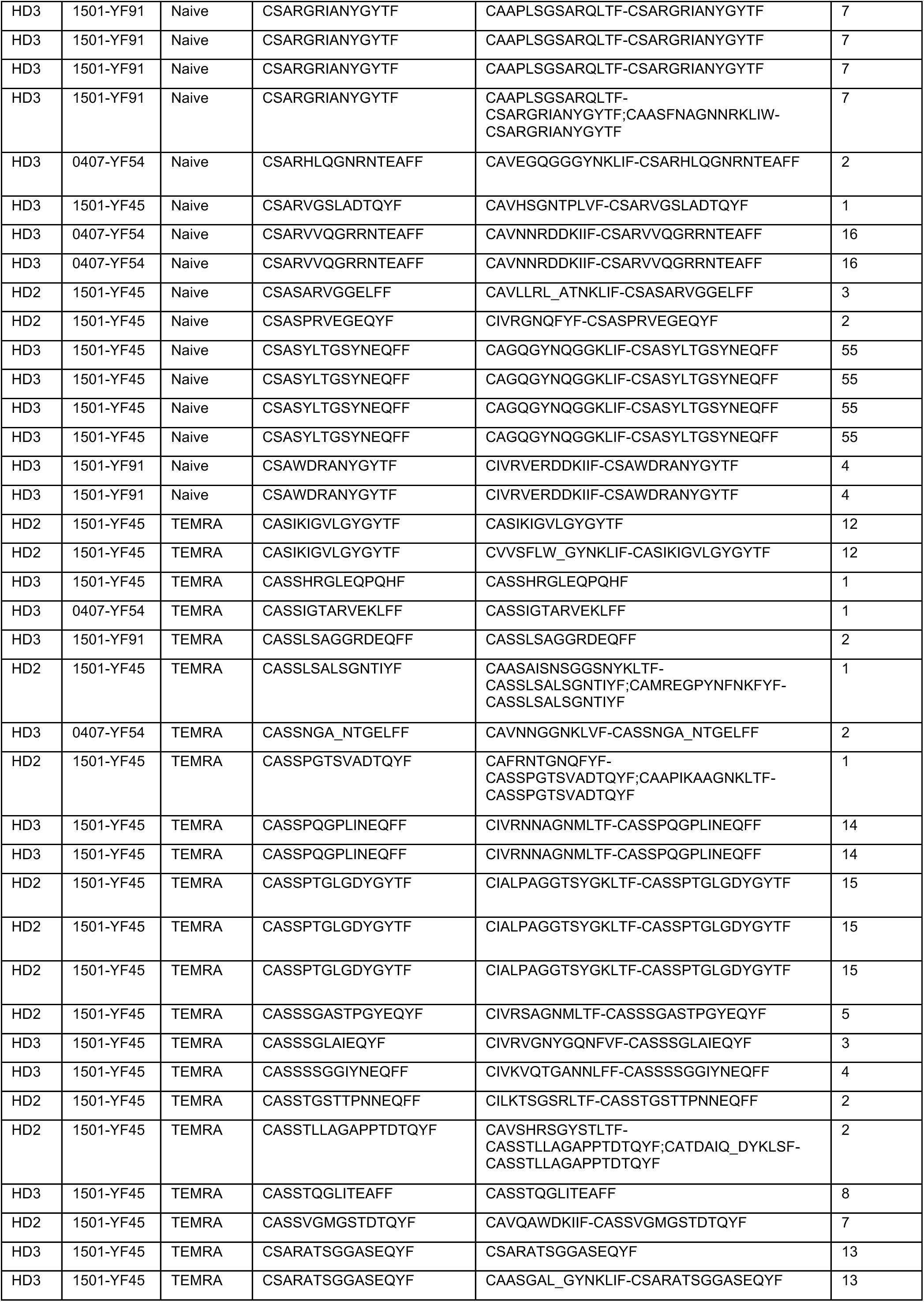
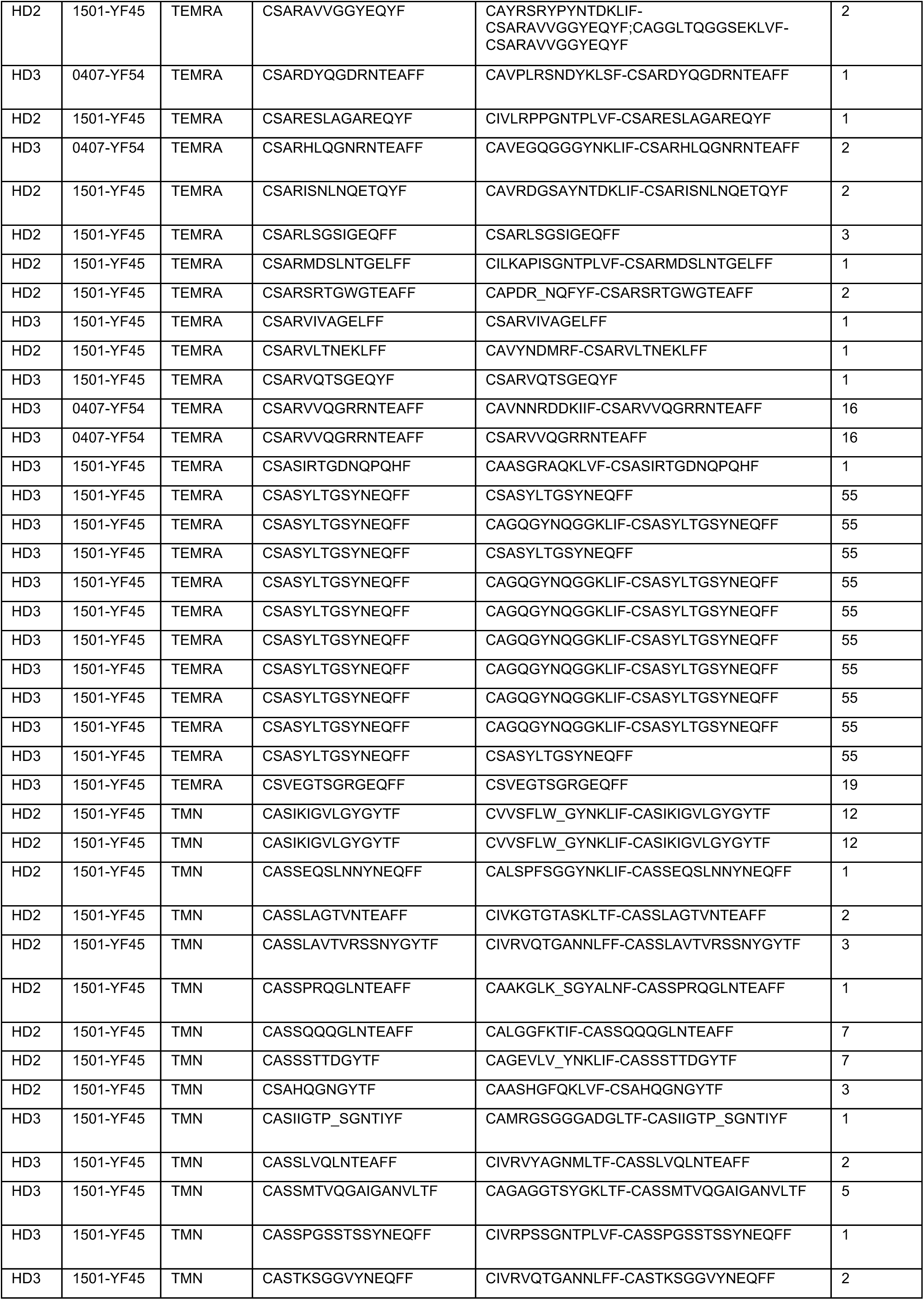
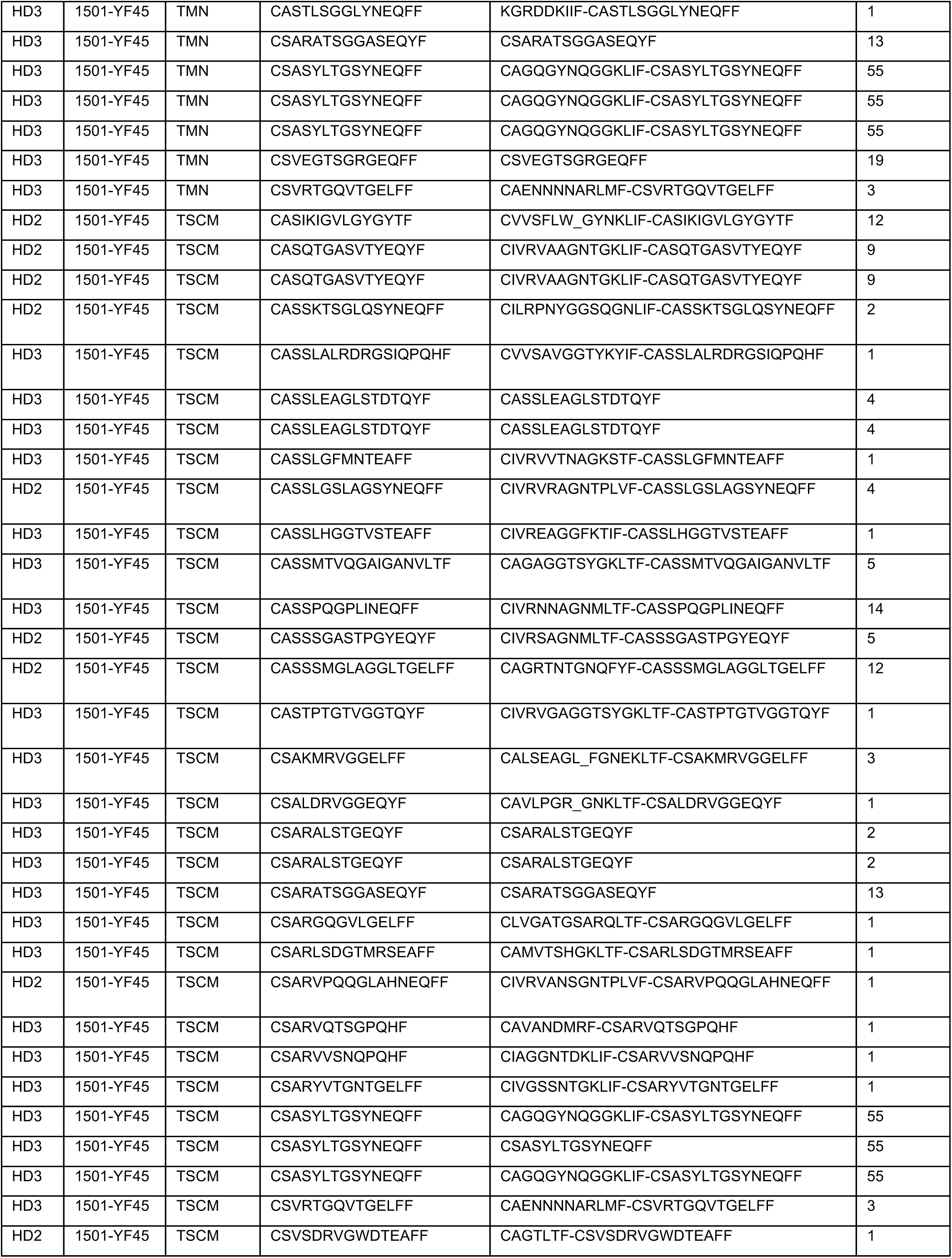
YFV-specific CD4^+^ T cells 7months or longer after vaccination

**Table S5:**
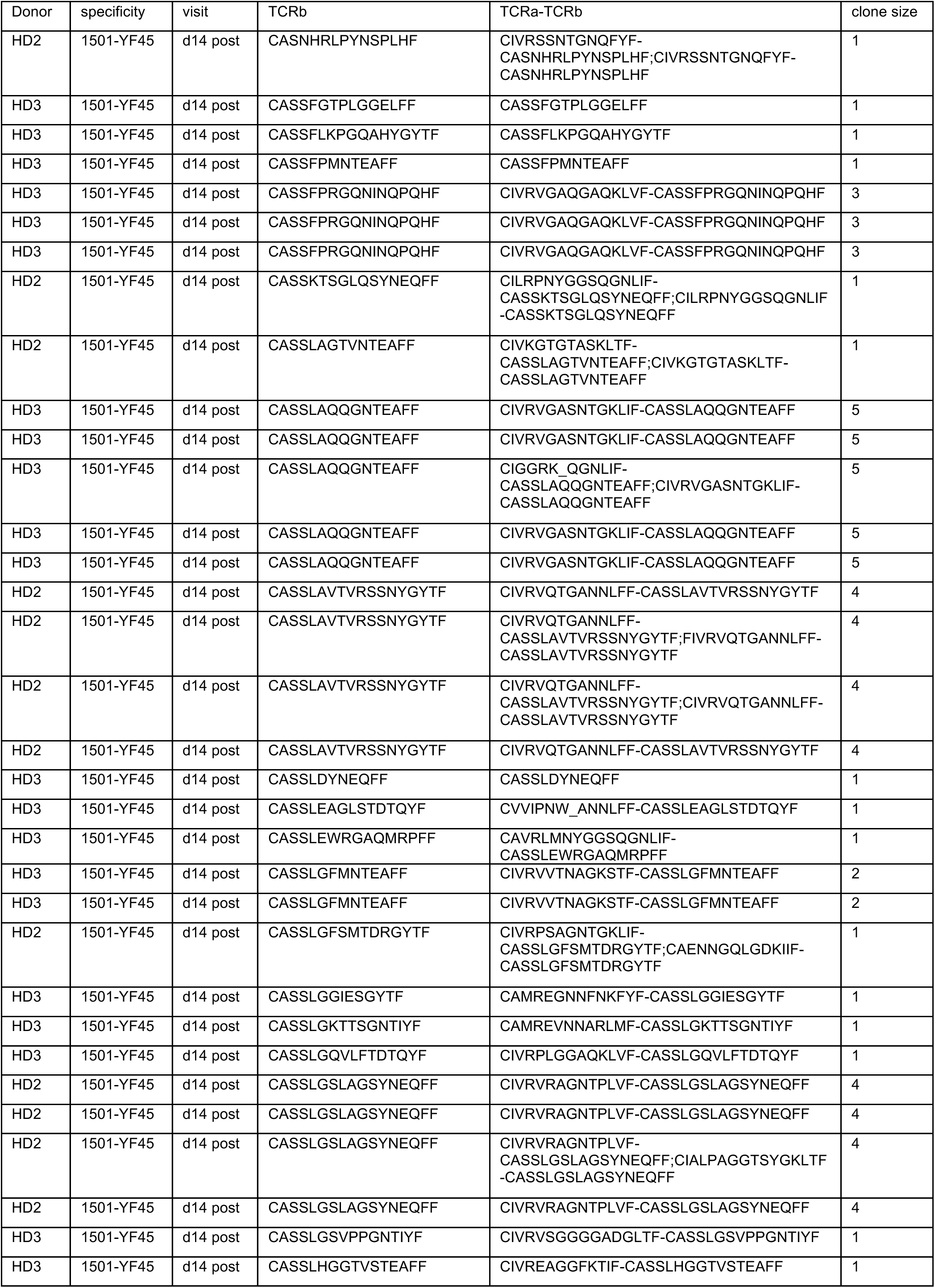
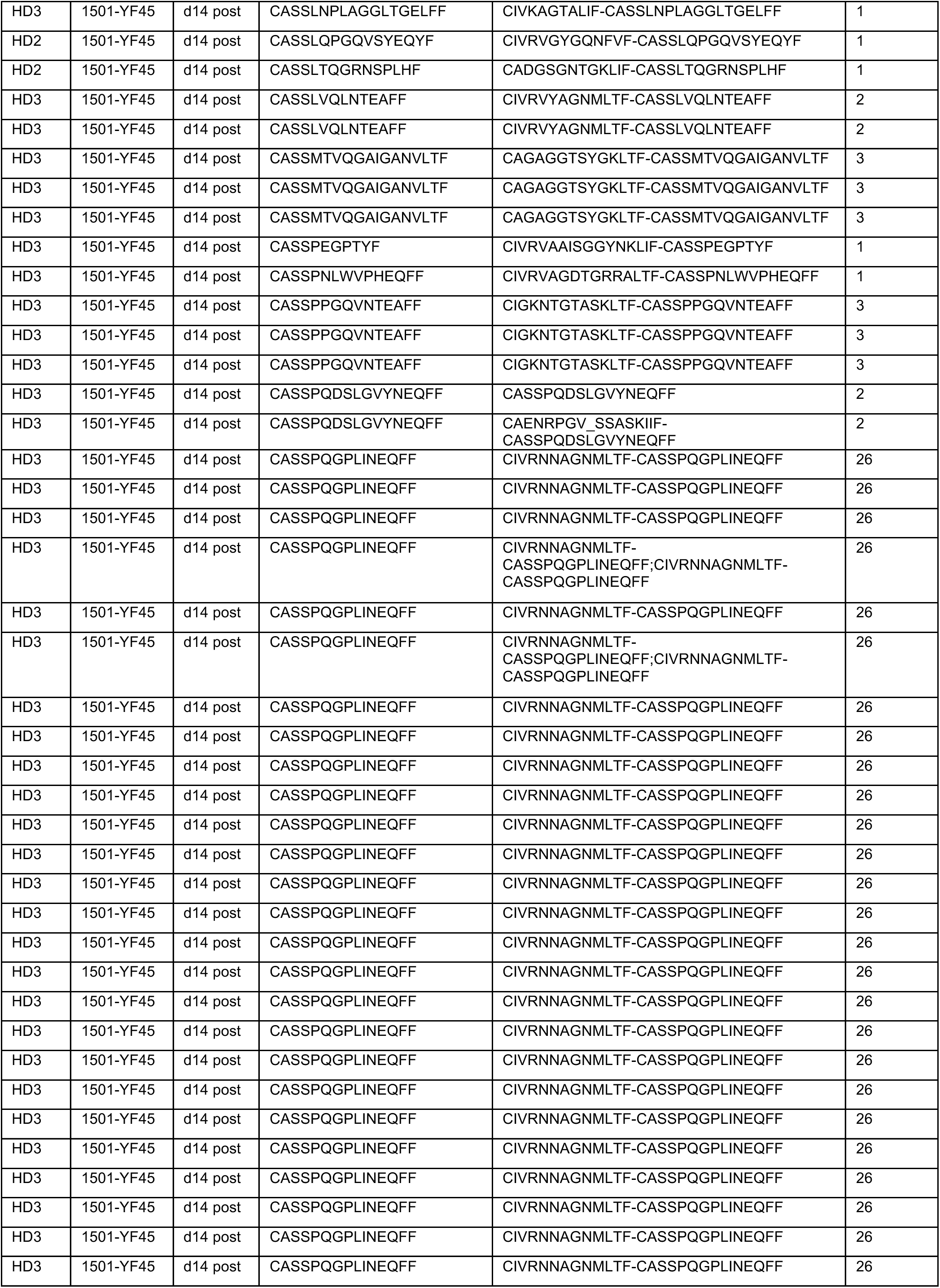
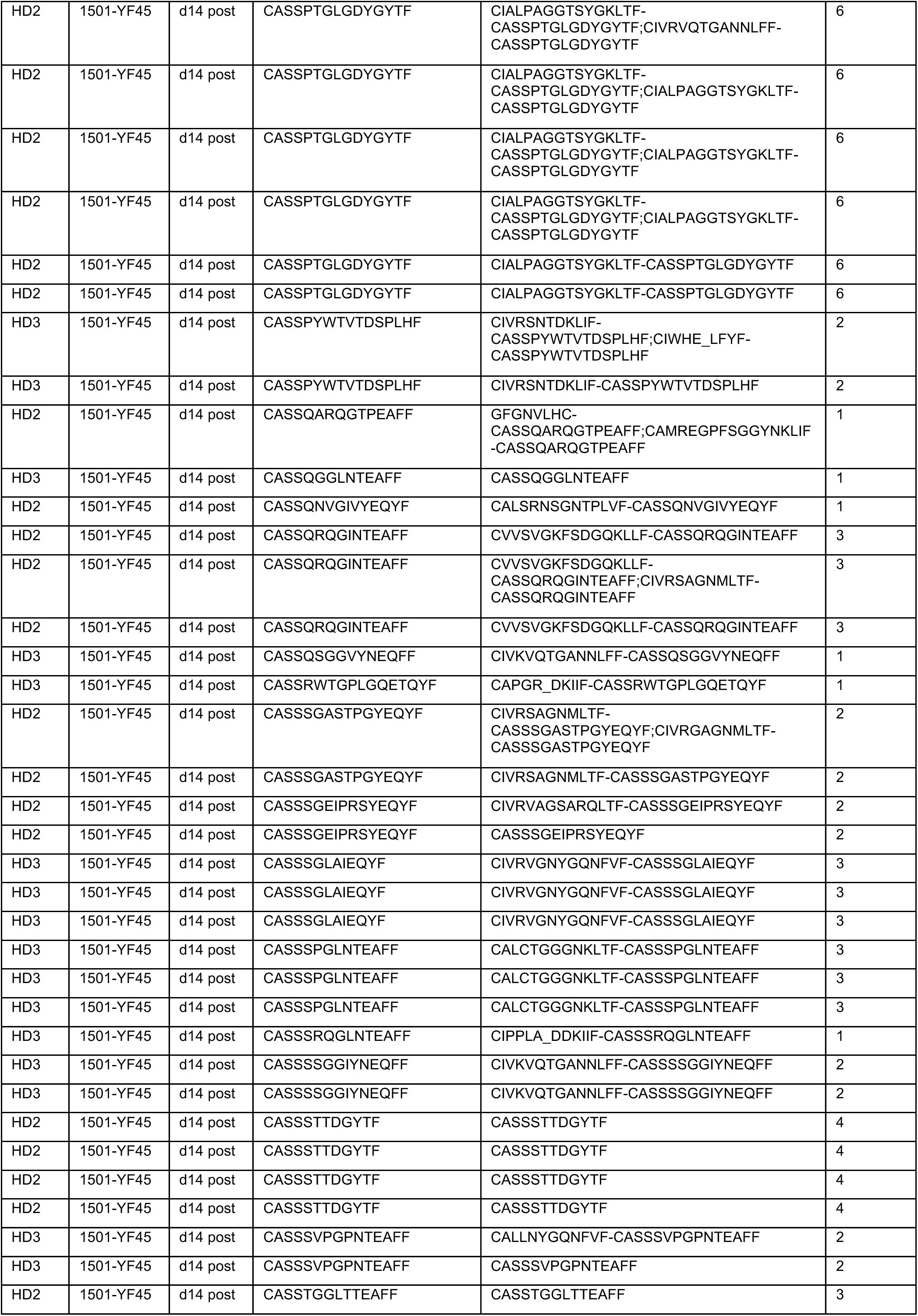
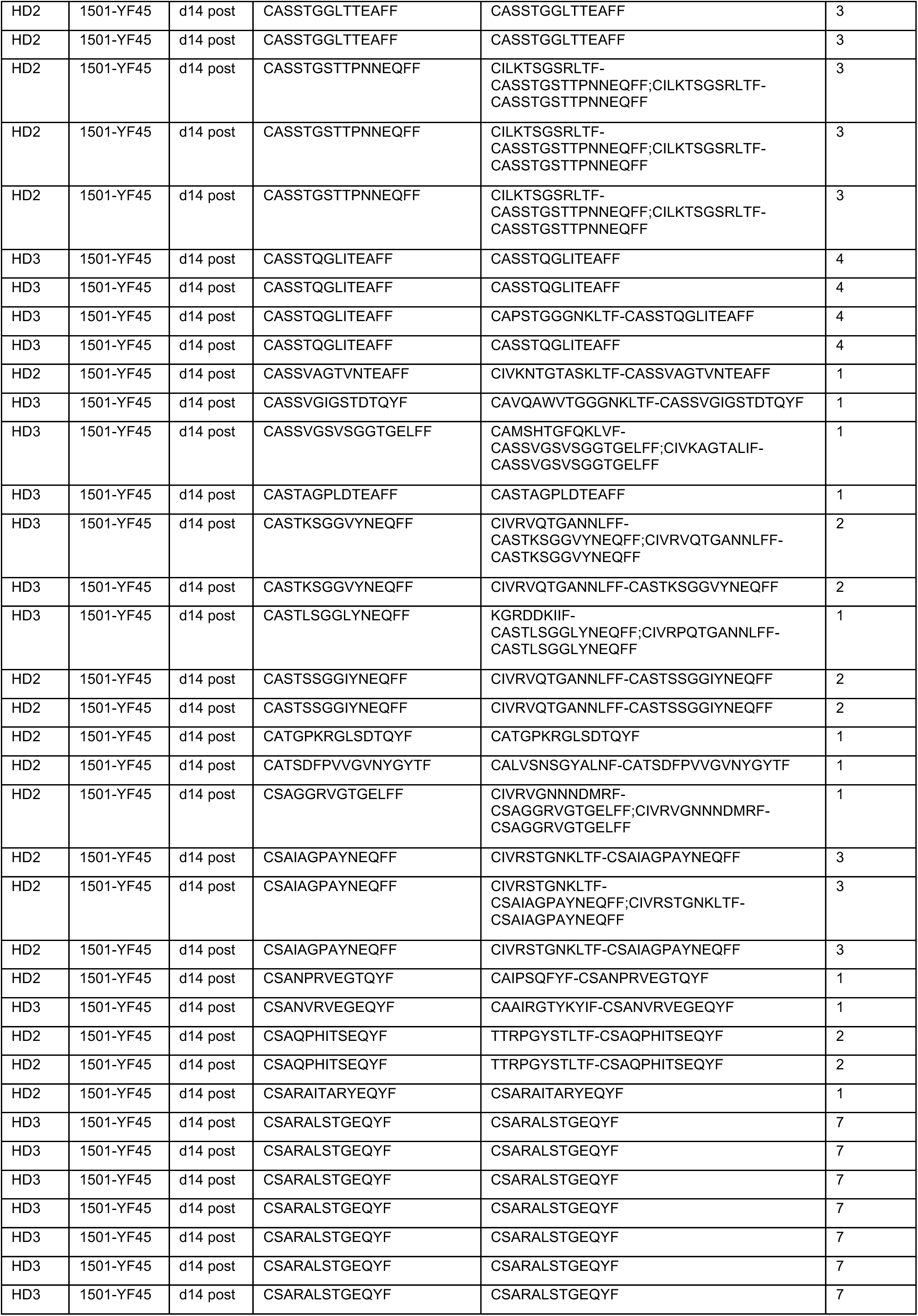
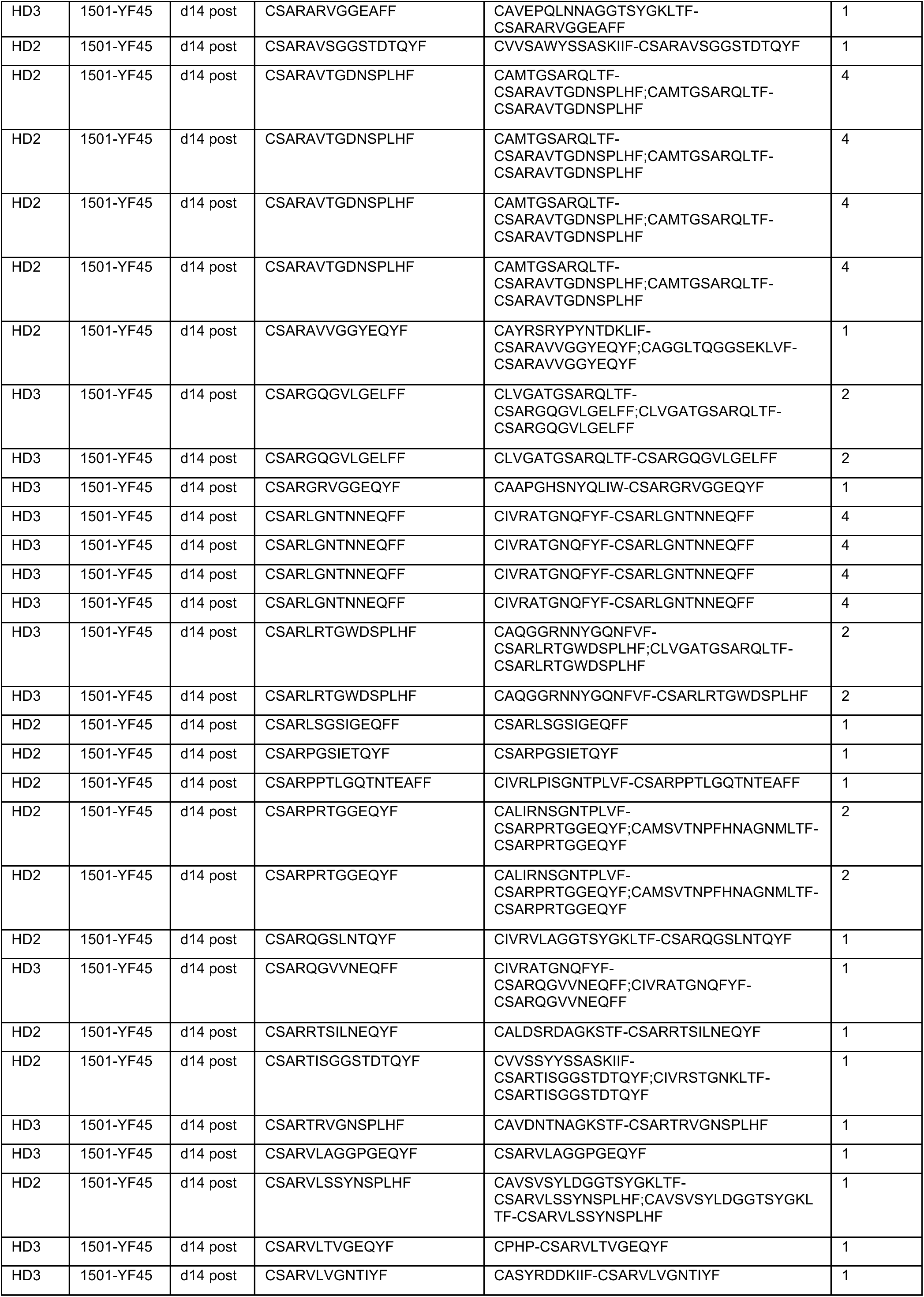
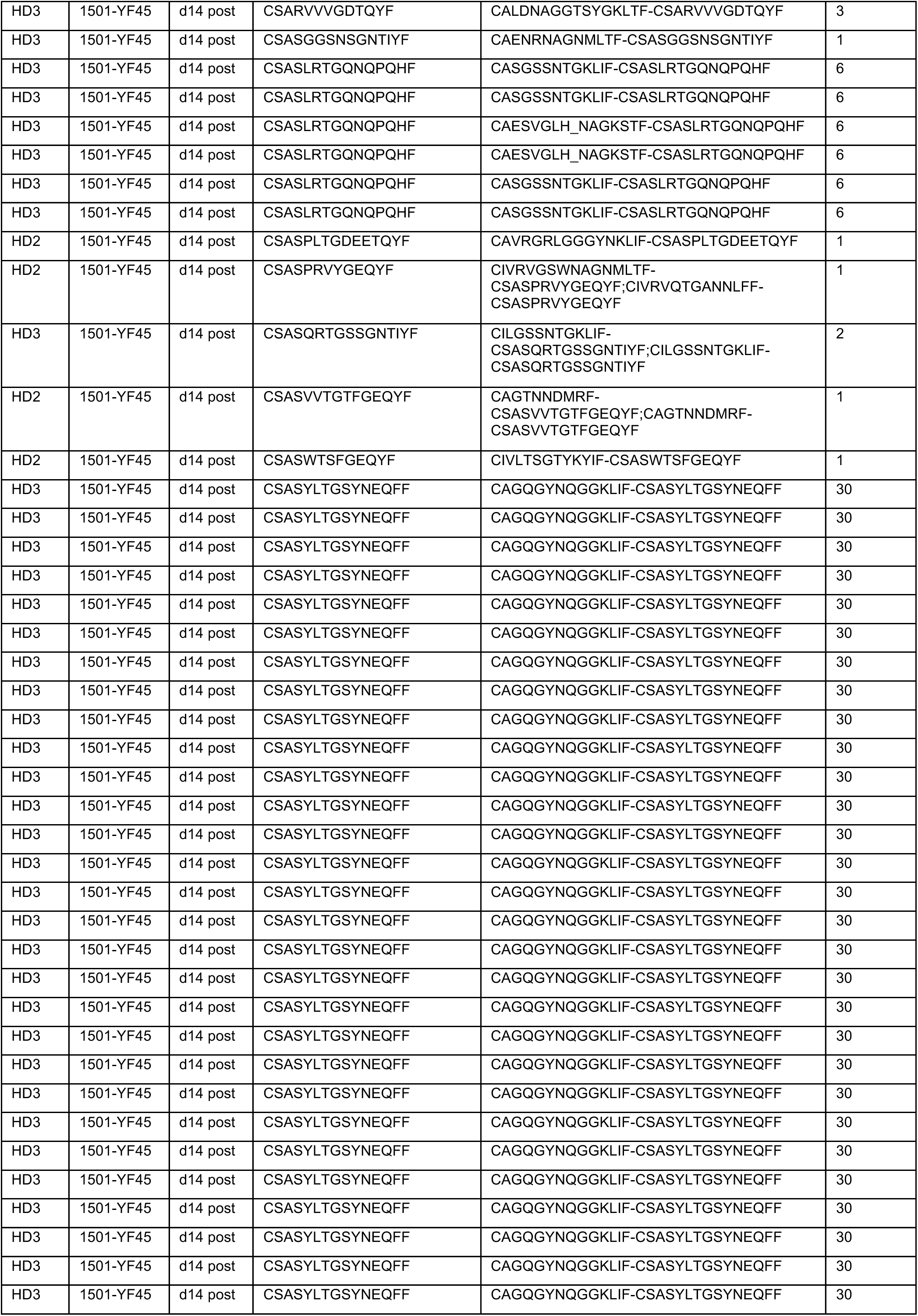
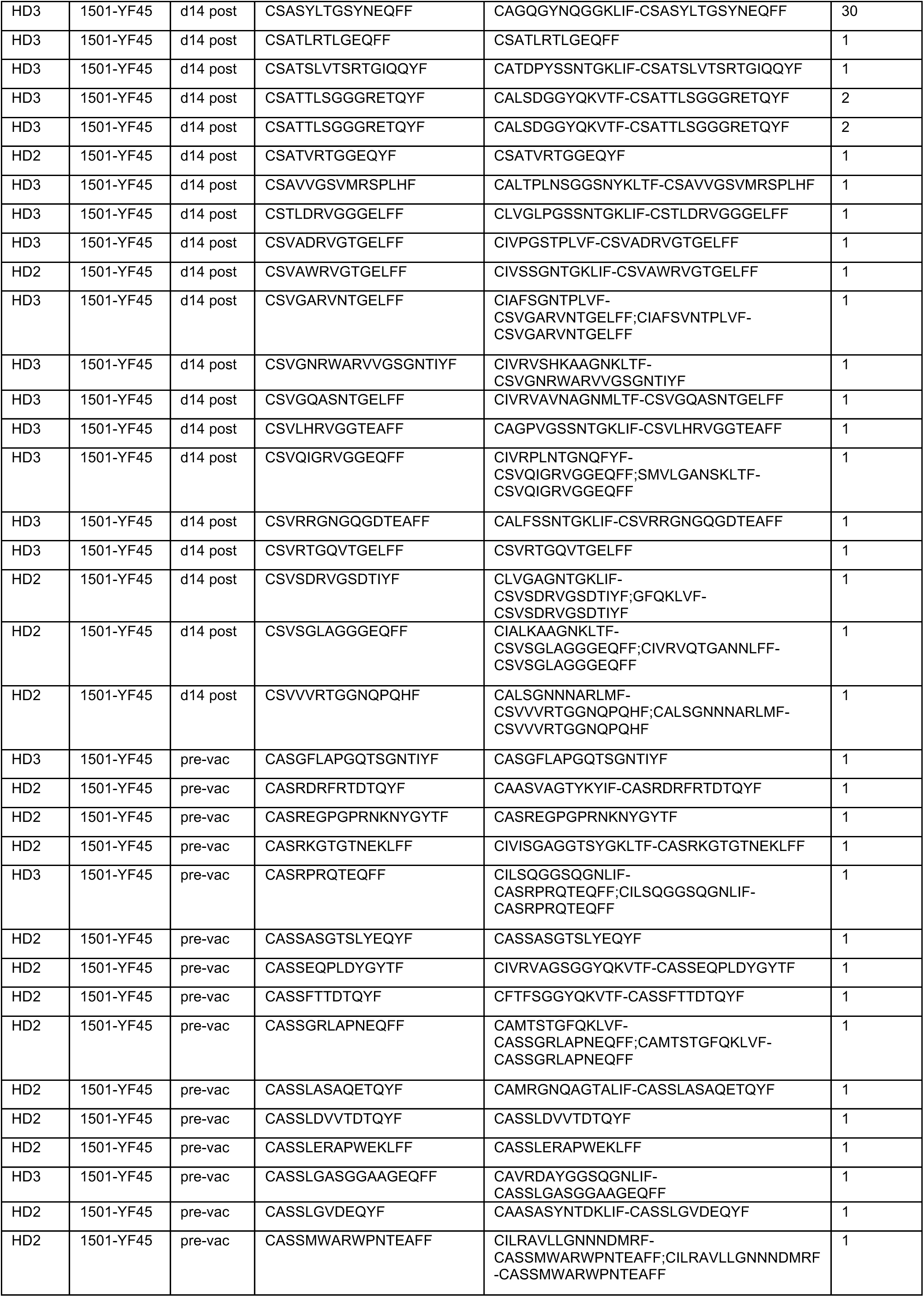
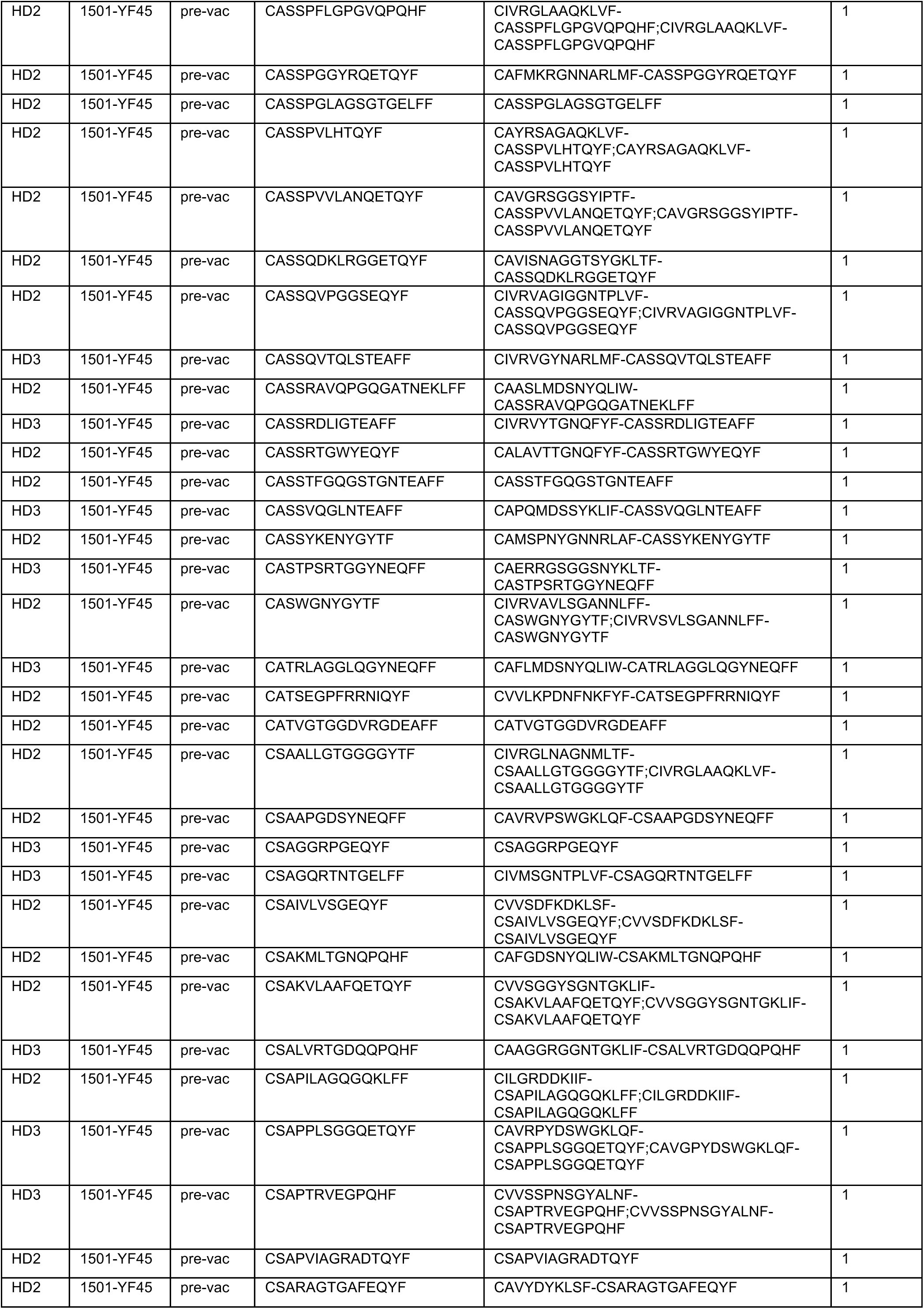
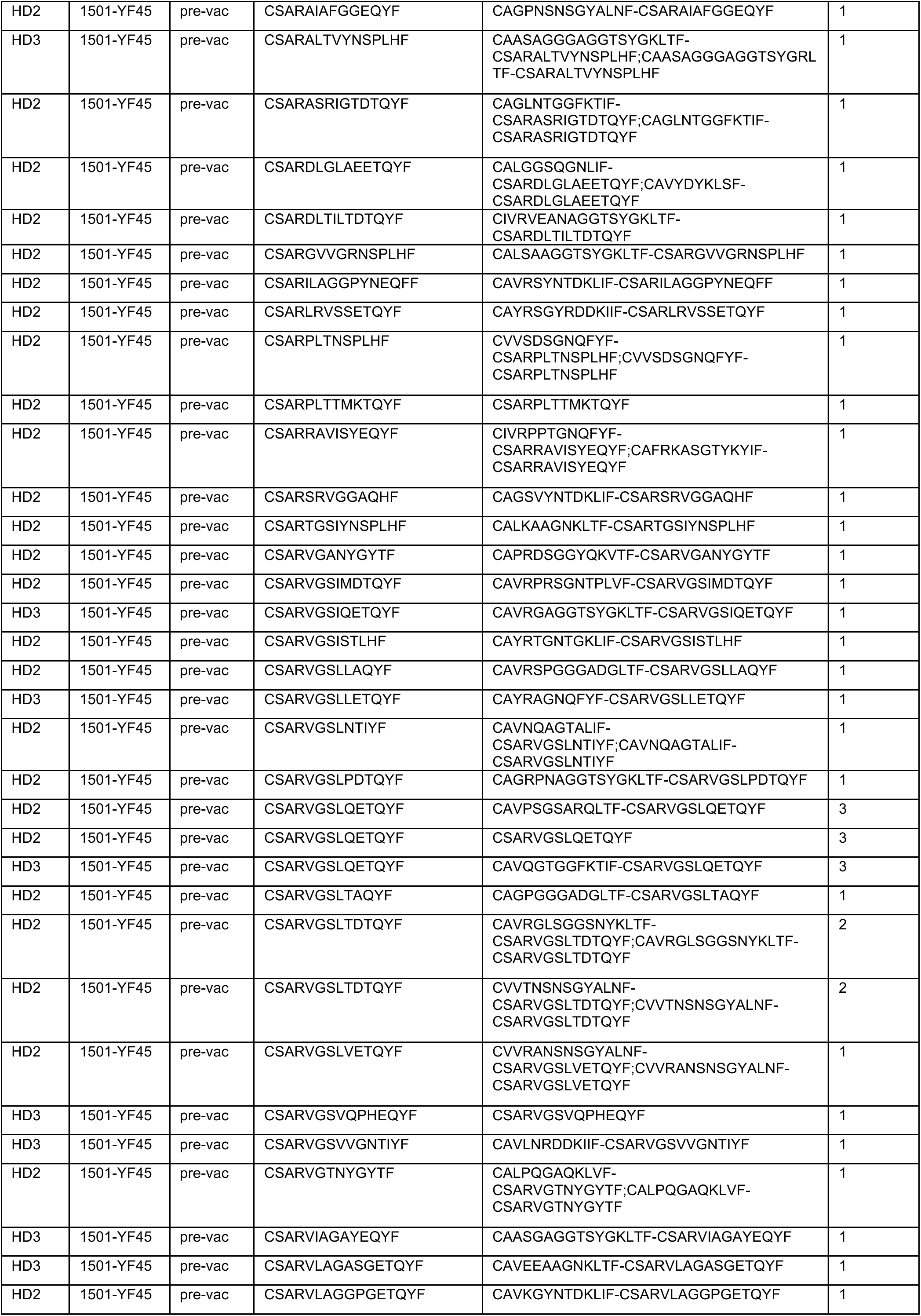
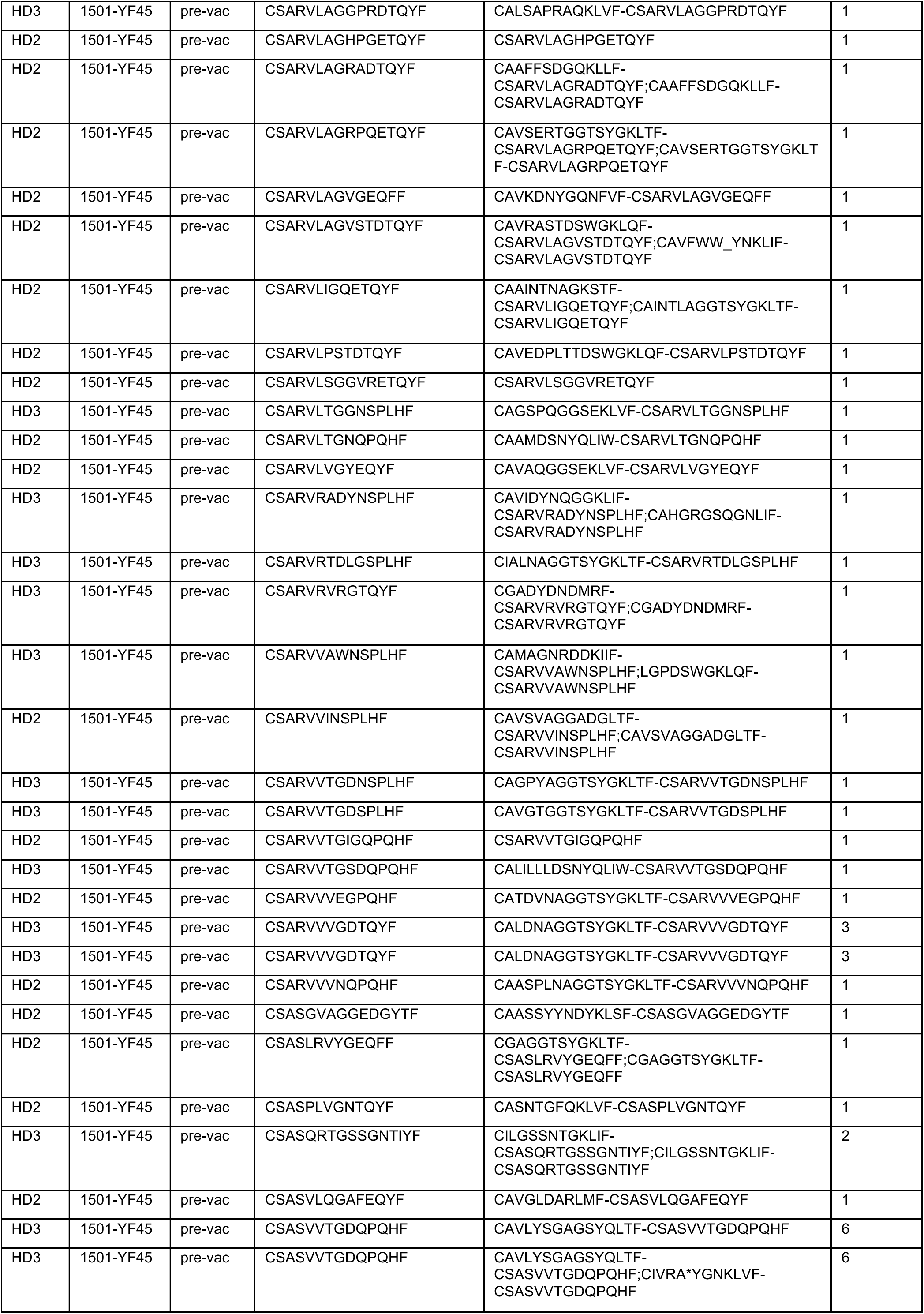
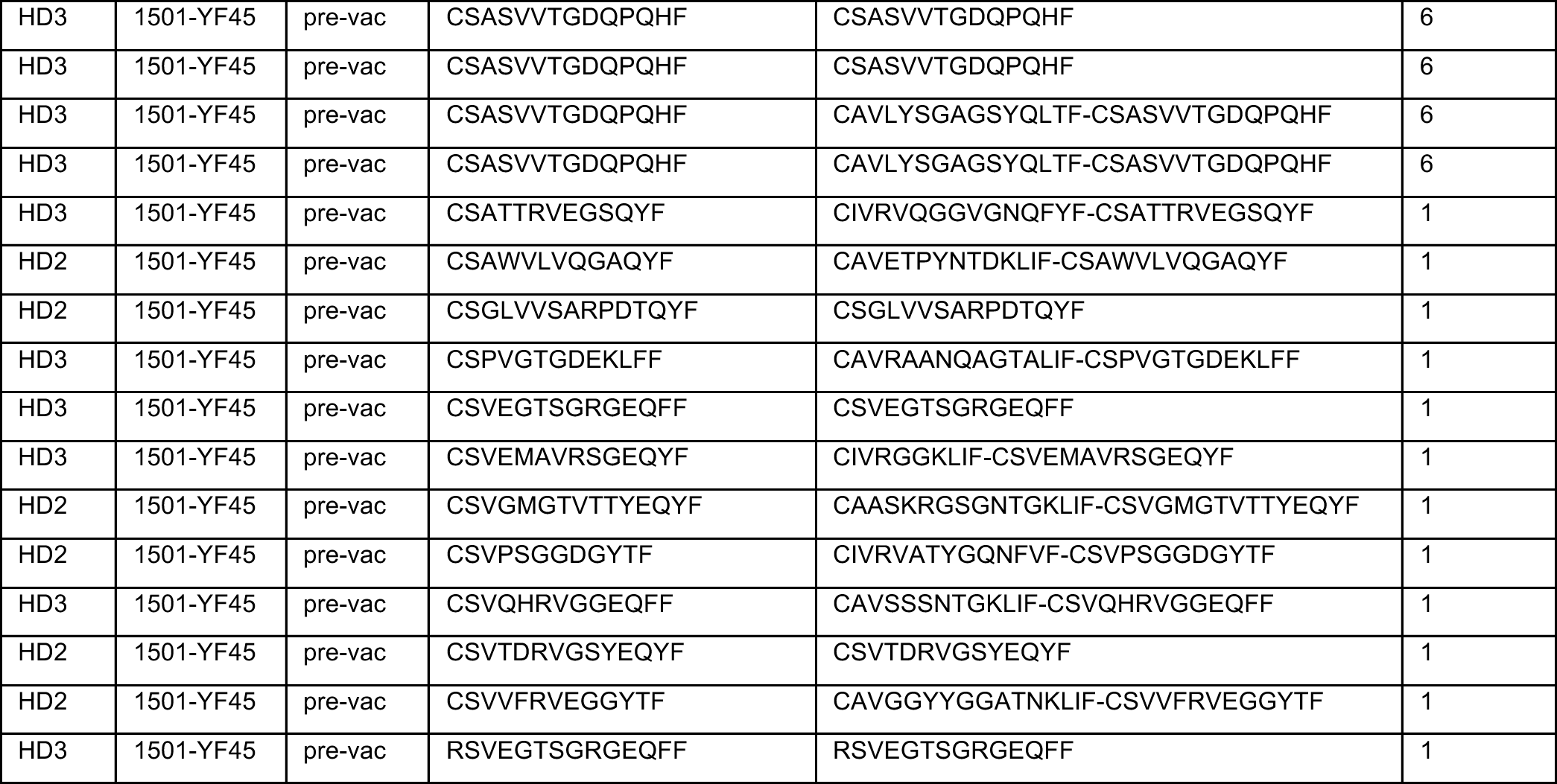
YF45 tetramer^+^ T cells before and 14 days after YFV vaccination

**Table S6:**
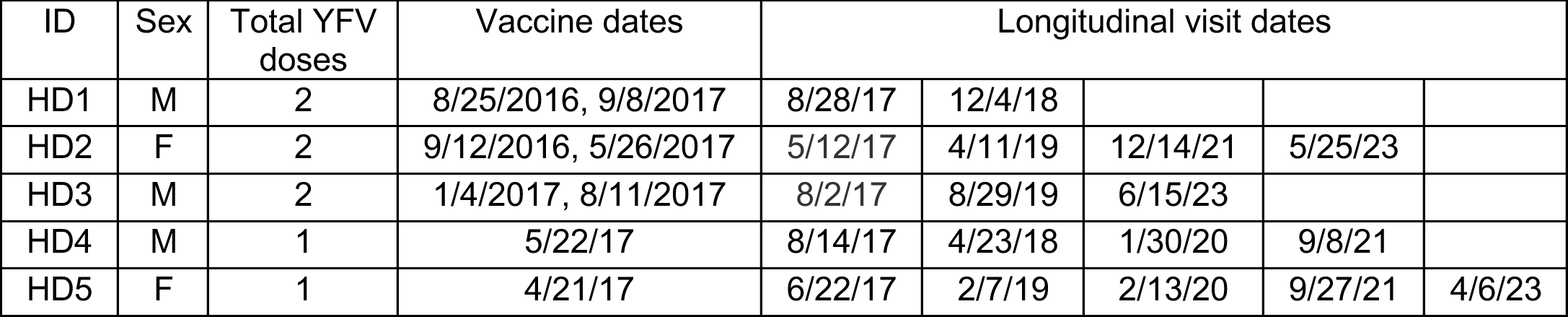
Longitudinal follow-up visits

| ID | Sex | Total YFV doses | Vaccine dates | Longitudinal visit dates |  |  |  |  |
| --- | --- | --- | --- | --- | --- | --- | --- | --- |
| HD1 | M | 2 | 8/25/2016, 9/8/2017 | 8/28/17 | 12/4/18 |  |  |  |
| HD2 | F | 2 | 9/12/2016, 5/26/2017 | 5/12/17 | 4/11/19 | 12/14/21 | 5/25/23 |  |
| HD3 | M | 2 | 1/4/2017, 8/11/2017 | 8/2/17 | 8/29/19 | 6/15/23 |  |  |
| HD4 | M | 1 | 5/22/17 | 8/14/17 | 4/23/18 | 1/30/20 | 9/8/21 |  |
| HD5 | F | 1 | 4/21/17 | 6/22/17 | 2/7/19 | 2/13/20 | 9/27/21 | 4/6/23 |

## Acknowledgments

We thank our study subjects for their participation.

## Author contributions

Conceptualization, L.F.S.; Experimentation, Y.P.; Sequence analyses, L.B.; High-dimensional phenotypic analyses, R. X.; Study recruitment, B.P; Modeling and statistical support, V.Z.; Supervision, L.F.S.; Manuscript preparation, L.F.S., L.B., Y.P, and V. Z.

## Data and material availability

All data needed to evaluate the conclusions in the paper are present in the paper or the supplementary materials.

